# Reconstructing the 3D genome organization of Neanderthals reveals that chromatin folding shaped phenotypic and sequence divergence

**DOI:** 10.1101/2022.02.07.479462

**Authors:** Evonne McArthur, David C. Rinker, Yang Cheng, Qixuan Wang, Juan Wang, Erin N. Gilbertson, Geoff Fudenberg, Maureen Pittman, Kathleen Keough, Feng Yue, Katherine S. Pollard, John A. Capra

## Abstract

Changes in gene regulation were a major driver of the divergence of archaic hominins (AHs)—Neanderthals and Denisovans—and modern humans (MHs). The three-dimensional (3D) folding of the genome is critical for regulating gene expression; however, its role in recent human evolution has not been explored because the degradation of ancient samples does not permit experimental determination of AH 3D genome folding. To fill this gap, we apply novel deep learning methods for inferring 3D genome organization from DNA sequence to Neanderthal, Denisovan, and diverse MH genomes. Using the resulting 3D contact maps across the genome, we identify 167 distinct regions with diverged 3D genome organization between AHs and MHs. We show that these 3D-diverged loci are enriched for genes related to the function and morphology of the eye, supra-orbital ridges, hair, lungs, immune response, and cognition. Despite these specific diverged loci, the 3D genome of AHs and MHs is more similar than expected based on sequence divergence, suggesting that the pressure to maintain 3D genome organization constrained hominin sequence evolution. We also find that 3D genome organization constrained the landscape of AH ancestry in MHs today: regions more tolerant of 3D variation are enriched for introgression in modern Eurasians. Finally, we identify loci where modern Eurasians have inherited novel 3D genome folding patterns from AH ancestors and validate folding differences in a high-frequency locus using Hi-C, revealing a putative molecular mechanism for phenotypes associated with archaic introgression. In summary, our application of deep learning to predict archaic 3D genome organization illustrates the potential of inferring molecular phenotypes from ancient DNA to reveal previously unobservable biological differences.

## 1 Introduction

The sequencing of archaic hominin (AH) and modern human (MH) genomes has transformed our understanding of human history, evolution, and biology [1–5]. However, even with these whole-genome sequences available, our understanding of how and why AHs differed from MHs is limited [6]. A major challenge in understanding the phenotypic and sequence differences between AHs and MHs is bridging the gap between genetic variation and function. The evolution of hominins is largely driven by changes in the regulation of conserved proteins [7–13], but the mechanisms through which archaic variants influence gene expression, and ultimately phenotype, are incompletely understood [6, 13, 14].

Many studies that investigate the gene regulatory differences between MHs and AHs leverage Neanderthal ancestry remaining in modern Eurasians. Because MHs interbred with many AH groups over the past 50,000 years, more than one-third of the Neanderthal genome remains in introgressed sequences in MH genomes [15, 16]. These investigations have found widespread expression differences between Neanderthal and MH alleles [11, 12], many of which are hypothesized to contribute to trait variation in diverse MHs [17–21]. Phenotypes associated with Neanderthal ancestry range from immune system response [18, 19, 22–29], hair and skin coloration [18, 19, 30–32], metabolism [33–36], cardiopulmonary function [19, 37], skeletal morphology [19, 38], and behavior [18, 19]. However, since most regions of MH genomes have little or no evidence of introgression [11, 12, 30, 31, 39–41], considering only introgressed variation provides a very limited view into hominin biology and cannot address why certain regions of MH genomes tolerated Neanderthal DNA better than others.

Colbran et al. [13] addressed this challenge by inferring AH gene regulation genome-wide through predictive models trained on gene expression data in MHs [42]. They estimated that over 1900 genes had different patterns of regulation between AHs and MHs. However, the specific molecular mechanisms through which archaic variants alter gene expression remain unclear. Gokhman et al. [43] and Batyrev et al. [44] aimed to elucidate these mechanisms by computationally reconstructing maps of AH DNA methylation. They found 2,000 differentially methylated regions that associate with genes predominantly related to facial and limb anatomy. Together, these illustrate the potential to mechanistically link archaic genotypes with regulatory functions via prediction of molecular phenotypes.

Yet, previous work has been unable to address a fundamental aspect of gene regulation and genome function—the physical three-dimensional (3D) organization of the genome. Regulation of gene expression is facilitated by the 3D looping and folding of chromatin in the cell nucleus, which is central to enhancer-promoter (E-P) communication and insulation [45–52]. The 3D genome also plays a role in determining cell-type identity, cellular differentiation, replication timing, and risk for multiple diseases [53– 59]. Advances in chromosome-conformation-capture technologies (3C, 4C, 5C, Hi-C, MicroC) [60–64] allow quantification of genome folding at increasing resolution from chromosomal territories, megabase-scale topologically associating domains (TADs), to smaller-scale loops [62] and “architectural stripes,” which can reflect enhancer activity and gene activation [65–67]. Disrupting 3D genome folding can cause inappropriate E-P interactions and alter gene expression in ways that lead to disease [49, 50, 68–72]. Accordingly, there is preliminary evidence suggesting the 3D genome constrains variation at different scales of evolution [73–77] and that reorganization of chromatin may contribute to gene regulatory evolution and inter-species gene expression divergence [78].

Thus, to fully understand the consequences of genetic variation between AHs and MHs, we must consider the 3D genome folding. However, the role of 3D genome organization in the divergence between AHs and MHs has never been explored because chromatin contacts cannot be assayed in ancient DNA. 3D genome folding is facilitated by a complex interplay of CTCF binding with cohesin and other architectural factors [50, 62, 79, 80]. Recent deep learning methods have been developed that learn the sequence “grammar” underlying 3d genome folding patterns [81–84]. We hypothesized that these deep learning methods would allow us to infer genome-wide 3D chromatin contact maps of Neanderthals and Denisovans. Because the molecular mechanisms that determine genome organization, like CTCF binding and co-localization with cohesin, are largely evolutionarily conserved [85, 86], models trained using human data perform well even when applied to DNA sequences from distantly related species, such as mouse [82]. Thus, unlike genome-wide methods for predicting organism-level phenotype (e.g., polygenic risk scores), these models can be applied across diverse hominins.

To elucidate the contribution of 3D genome folding to recent hominin evolution, we apply novel deep learning methods for inferring 3D genome organization from DNA sequence patterns to Neanderthal, Denisovan, and diverse MH genomes. Using the resulting genome-wide 3D genome folding maps, we identify 167 loci that are divergent in 3D organization between AHs and MHs. We show that these 3D-diverged loci are enriched for physical links to genes related to the function and morphology of the eye, supra-orbital ridge, hair, lung function, immune response, and cognition. We also find that 3D genome organization constrained recent human evolution and patterns of introgression. Finally, we evaluate the legacy of introgression on the 3D organization of humans and identify examples where introgression imparted divergent 3D genome folding to Eurasians. In summary, our application of deep learning to predict archaic 3D genome folding provides a window into previously unobservable molecular mechanisms linking genetic differences to phenotypic consequences in hominin evolution.

## 2 Results

### 2.1 Reconstructing the 3D genome organization of archaic hominins

To evaluate the role of 3D genome organization changes in recent human evolution, we apply deep learning to infer 3D genome organization from DNA sequences of archaic hominins (AHs) and modern humans (MHs) (Fig. 1). We consider the genomes of four AHs—one Denisovan and three Neanderthals, each named for where they were discovered (*Altai* mountains, *Vindija* and *Chagyrskaya caves*) [1–4]. We compare these to 20 diverse MHs from the 1000 Genomes Project (Table S1) [87].

**Figure 1:**
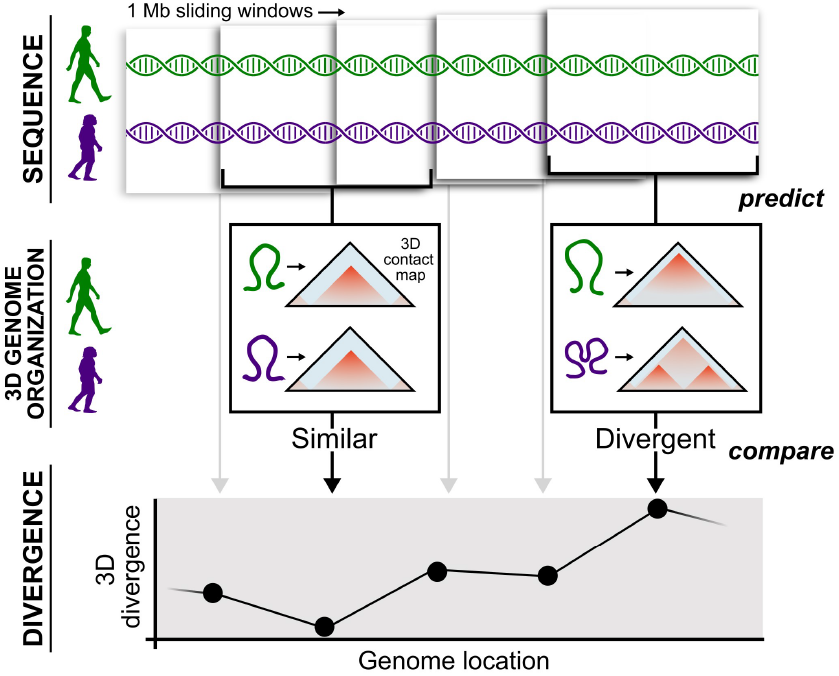
Reconstructing the 3D genome organization of archaic hominins. We infer 3D genome organization from sequence across the genomes of modern humans (MHs, green) and archaic hominins (AHs, purple). Using approximately 1 Mb (1,048,576 bp) sliding windows (overlapping by half), we input the genome sequences into Akita, a convolutional neural network, to predict 3D genome contact maps [82]. The resulting contact maps are compared between MHs and AHs to identify regions that have similar 3D genome organization (left, low divergence) and regions that have different 3D organization (right, high divergence).

For each individual, we predict chromatin contact maps across the genome. Each contact map gives a 2D representation of the predicted 3D chromatin physical contacts, which will refer to as “3D genome organization”. We predict these maps using approximately 1 Mb (1,048,576 bp) tiled sliding windows overlapping by half with Akita, a convolutional neural network (CNN) trained on high-quality experimental chromatin contact maps (Hi-C and Micro-C) [82]. Each resulting contact map represents pairwise physical 3D contact frequencies at approximately 2 kb (2,048 bp) resolution for a single individual. Previous work demonstrated that Akita accurately infers 3D contact organization at this resolution [82]. We only consider windows with full (100%) sequence coverage in the MH reference, and we conservatively mask missing archaic sequence with the human reference sequence (Figs. S1,S2,S3 and Methods).

We compare contact maps from two genomes using a “3D divergence” score, namely, one minus the Spearman’s rank correlation coefficient (1 − *ρ*) for all pixels in the maps. Genomic windows with more different 3D genome maps have higher 3D divergence and, conversely, a window with lower 3D divergence will reflect more 3D similarity (Fig. 1). Other divergence metrics (e.g., based on Pearson’s correlation coefficient and mean squared difference) are strongly correlated (Fig. S4) and our previous work has shown that simple correlation-based metrics are an appropriate method of genome-wide prioritization for 3D comparisons when compared to more complex computationally-intensive methods [88]. Akita is trained simultaneously on Hi-C and Micro-C across five cell types in a multi-task framework. In the main text we focus on predictions from the highest resolution cell type, human foreskin fibroblast (HFF). Results are similar when considering other cell types (e.g. embryonic stem cells) (Fig. S5), likely because of limited cell-type-specific differences in both available experimental data and model predictions [82].

### 2.2 Archaic hominin and modern human genomes exhibit a range of 3D divergence

Reconstructing the genome-wide 3D genome organization of AHs and MHs revealed genomic windows with a range of 3D divergence (Fig. 2A). Most of the genome has very similar 3D genome organization between AHs and MHs (circle example in Fig. 2A-B). However, we also found regions of AH-MH 3D genome divergence. Some of these differences are changes in predicted chromatin contact intensity but similar overall organization (diamond example in Fig. 2A-B). Others reveal reorganization with evidence of new sub-organization (neo-TADs or -loops) or lost structures (fused TADs or loops) (indicated with an “x” example in Fig. 2A-B). At the 95^th^ percentile of observed divergence, differences in the contact maps are substantial. However, because the 3D divergence measure considers the entire window, strong focal changes may not rank as highly as structural differences that influence a large segment of the window (diamond vs. “x” examples in Fig. 2B).

**Figure 2:**
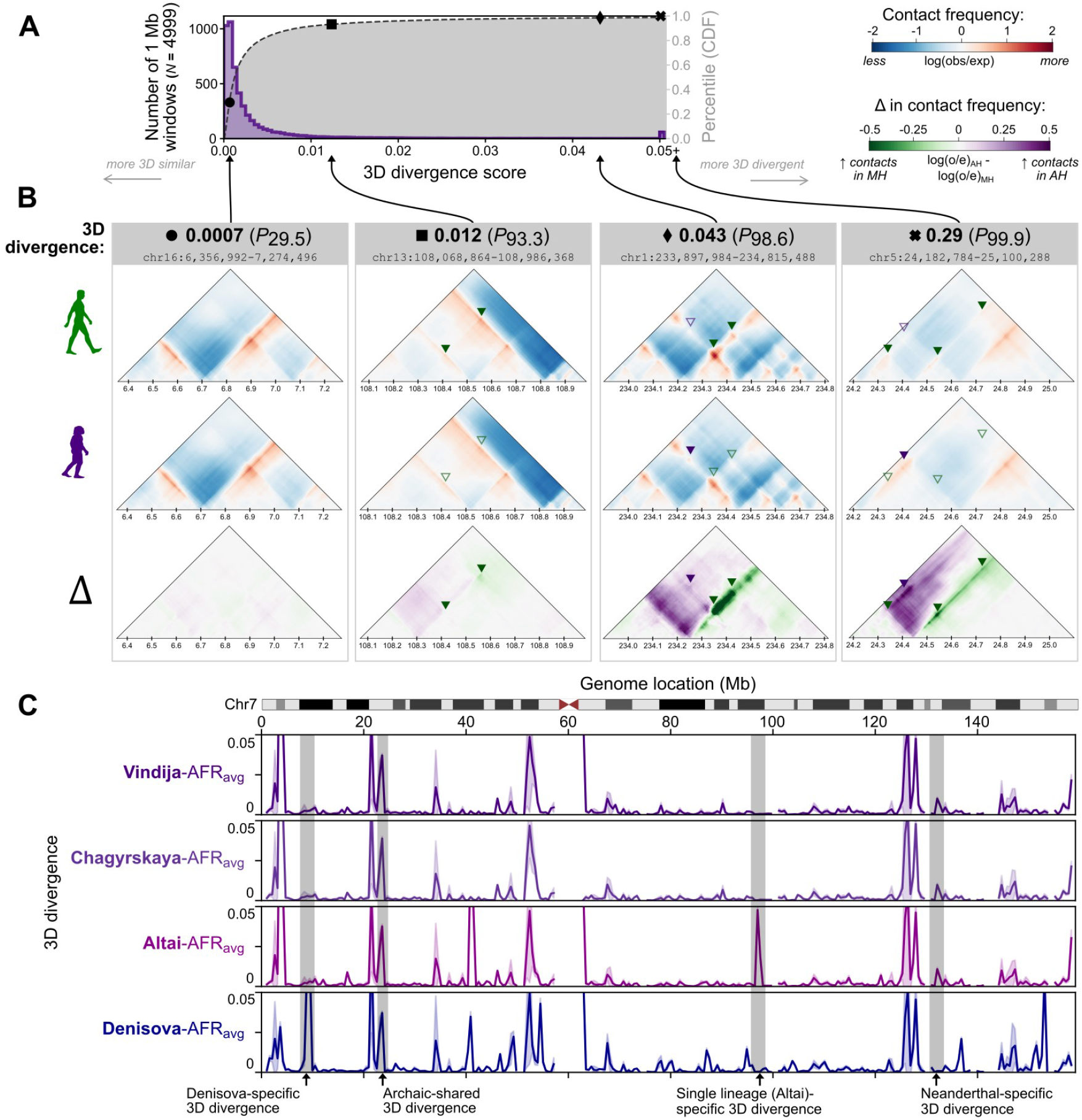
3D genome divergence between archaic hominins (AHs) and modern humans (MHs) varies across the genome. **(A)** Distribution of 3D genome divergence between AHs and modern humans MHs for 1 Mb windows across the genome. Most windows have similar 3D genome organization between MHs and AHs (low 3D divergence). The cumulative density function (CDF) of this distribution is overlaid in gray with percentiles on the right vertical axis. **(B)** We highlight four examples (shapes) along the 3D divergence distribution illustrating low 3D divergence (left) to high divergence (right). Each example compares a representative African MH (top, HG03105) to a Neanderthal (bottom, Vindija) in terms of both raw score (bolded number) and relative percentile of 3D divergence (number in parentheses). Examples with scores near the 95^th^ percentile have visible contact map differences, but the type of differences vary from re-organization (neo-TADs or TAD-fusions) to altered contact intensity (stronger vs. weaker TAD/loop). Green and purple triangles indicate regions with increased contact frequency in MH versus AH, respectively. (C) Average 3D divergence along chromosome 7 between AHs and five representative African MHs. The error band indicates the 95% confidence interval (CI). Comparing the 3D genomes of Neanderthals (purple) or Denisova (blue) with MHs reveals windows of both similarity and divergence (peaks). Featured examples (gray overlays) highlight regions of 3D divergence that are shared (e.g., shared across all archaics) or lineage-specific (e.g., specific to the Denisovan individual).

To illustrate genome-wide patterns of divergence in 3D organization, we plotted the average divergence of each of the AHs to five modern African individuals from different subpopulations (Fig. 2C). We show the landscape of 3D divergence across the entire genome for all four AHs in Fig. S6. Some AH-MH divergences are shared across all four archaics, while others are specific to a single lineage like the Denisovan individual (Fig. 2C). We only considered sub-Saharan Africans in these comparisons, because they have low levels of AH introgression. We consider how introgressed variation in Eurasians influences 3D divergence in a subsequent section.

### 2.3 3D genome organization diverges between AH and MH at 167 genomic loci

To consistently identify regions with divergent 3D genome organization between AH and MH, we compared the 3D contact maps at each locus for each AH to 20 MH (African) individuals. We applied a conservative procedure that required all 20 AH-MH comparisons to be more 3D divergent than all MH-MH comparisons (Fig. 3A). In other words, the differences between the 3D genome organization of an AH to all MHs must be more extreme than the differences between each MHs to all other MHs. Furthermore, we required the average AH-MH 3D divergence to be in the 80^th^ percentile of the most diverged. This identified regions with consistent 3D differences between AHs and MHs (Fig. 3A, left) while excluding regions with a large 3D diversity in modern humans (Fig. 3A, right) (Methods).

**Figure 3:**
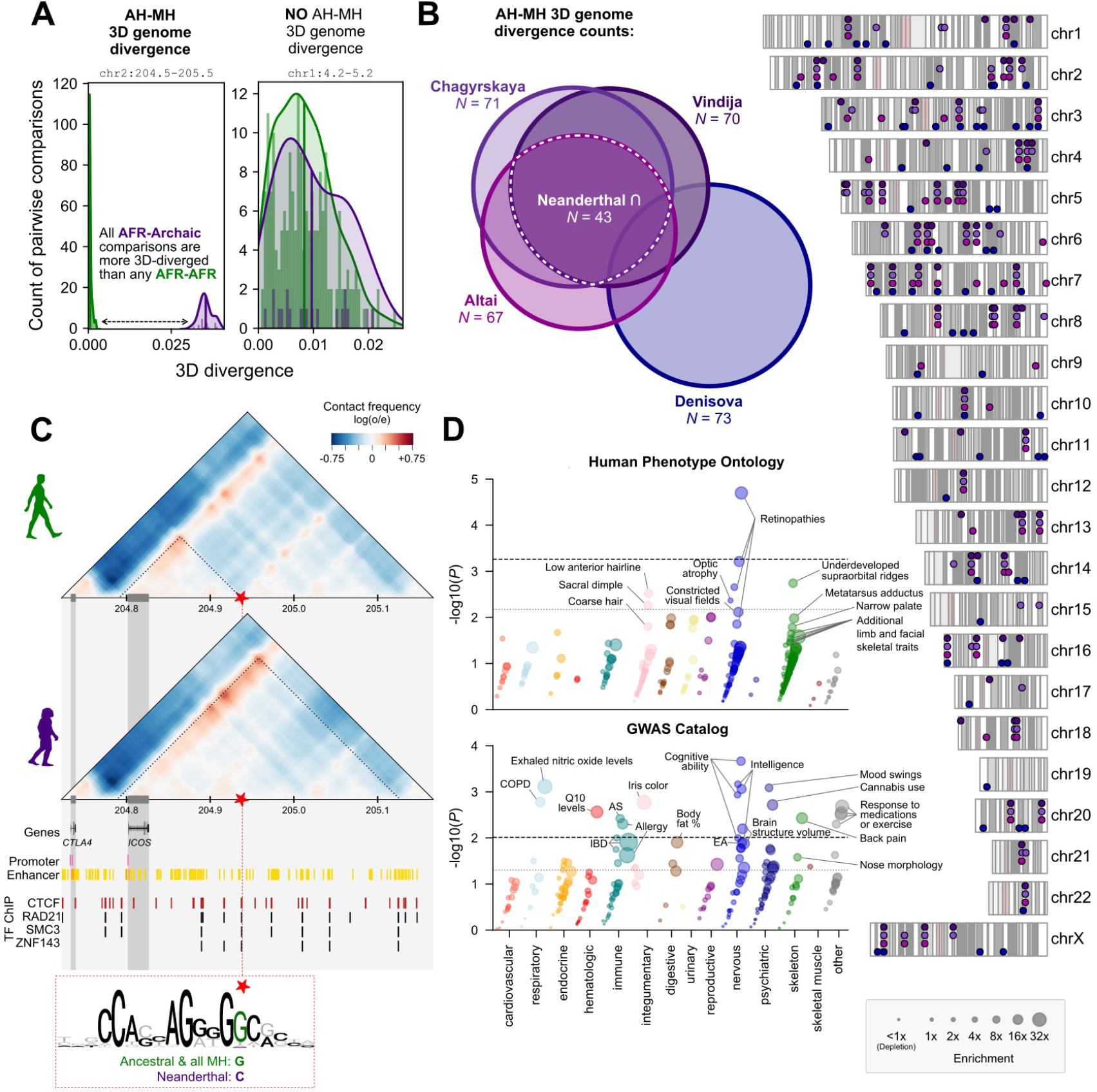
Regions with 3D divergence between MHs and AHs highlight loci linked to phenotypic differences. **(A)** We identified genomic windows with 3D divergence between AH and MH by comparing distributions of pairwise divergence in 3D contact maps. We used a conservative procedure that required all 20 comparisons of each AH to 20 MH (African) individuals (purple, *n* = 20) to be more 3D-diverged than all MH-MH comparisons (green, 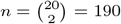) and the mean of the AH-MH divergences (purple) to be in the 80^th^ percentile of most diverged. The left plot shows an example that meets these criteria (chr2:204,472,320-205,520,896). The right shows an example where there is diversity in 3D genome organization, but not an AH-MH divergence (chr1:4,194,304-5,242,880). **(B)** We identified 167 AH-MH 3D divergent windows across the genome. Many are shared (Euler-diagram), but some are unique to a single lineage, with the most unique divergence in the Denisovan. **(C)** Contact maps for the example Neanderthal-MH 3D divergent window shown in **A** (zoomed to chr2:204,722,176-205,166,592). All MHs have a smaller domain insulated by a CTCF site (red star). In Neanderthals (Vindija and Altai), the CTCF motif is disrupted with a C instead of a G (red dashed box, chr2:204,937,347). We predict that this leads to ectopic connections with the promoter of *ICOS* (T-cell costimulator). (**D**) Phenotype enrichment for the 43 Neanderthal 3D diverged loci identified in **B** (white dashed line). We computed functional annotation enrichment for genes physically linked to 3D-modifying variants at these 3D divergent loci using HPO (top, *n* = 271) and GWAS catalog (bottom, *n* = 208) annotations (Methods). Within each phenotypic domain, traits are organized along the vertical axis by significance and along the horizontal axis by enrichment (also indicated by size). Genes nearby AH-MH 3D divergence are enriched for functions related to the retina and visual field, skeletal morphology (notably, supra-orbital ridge), hair, lung function, immune and medication response, and cognitive traits. Significance lines represent the *P* -value thresholds that controls the FDR with *q* = 0.05 (dotted) and *q* = 0.1 (dashed). (COPD: chronic obstructive pulmonary disease, AS: ankylosing spondylitis, IBD: inflammatory bowel disease, EA: educational attainment)

We find 167 total AH-MH consistently 3D diverged loci: 67, 70, 71, and 73 for Altai, Vindija, Chagyrskaya, and Denisova compared to MHs, respectively (Fig. 3B). 3D diverged loci are found through-out the genome on every chromosome (Fig. 3B). As suggested by Fig. 2C, some 3D divergences are shared by all four AHs (*N* = 7), and many are shared by all three Neanderthals (*N* = 43) (Fig. 3B). We summarize the AH-MH 3D divergent windows in Tables S2,S3 and report a larger set of windows based on less conservative criteria in Table S4. With this test, 3D divergent windows are extremely unlikely to be identified by chance. In 100,000 simulated genome-wide analyses with shuffled labels for the pairwise comparisons, we never observed a window meeting our criterion for 3D divergence (P ¡ 1×10^-5^). Moreover, when we computed the 3D divergent windows for each of the African individuals, we found a range of 10-30 divergent windows (Fig. S7).

To illustrate the properties of a AH-MH 3D divergent window, we highlight a divergent locus on chromosome 2 that is nearby several immune genes (Fig. 3C). MHs have an approximately 140 kb loop linking the promoter of *ICOS* at 204.80 Mb to a CTCF motif at 204.94 Mb. This CTCF motif is overlapped by many ChIP-seq peaks for transcription factors (TFs) involved in determining chromatin folding (CTCF, RAD21, SMC3, and ZNF143). The contact maps for both Vindija and Altai Neanderthal show a more prominent “architectural stripe”—an asymmetric loop-like contact often reflecting enhancer activity [65–67]—starting near the promoter of *ICOS*. However, in contrast to MHs, the loop does not end at the same CTCF site and instead has greater contact frequency with a CTCF site at 205.2 Mb. Thus, the resulting loop in Neanderthals is predicted to be over 400 kb—three times as large as the MH loop.

To determine which AH-MH nucleotide differences cause the largest change in the contact maps, we used *in silico* mutagenesis (Methods). Using an African MH (HG03105) background, we inserted every allele unique to the AH genome one-by-one and measured the resulting 3D genome divergence. This identifies the archaic variant resulting in the largest 3D organization changes between the AH and MH genomes, a G to C change at chr2:204,937,347 (Methods). This change disrupts a high information-content site in the CTCF binding site described above. All MHs carry an ancestral G allele, but Vindija and Altai have a derived C allele. In summary, we predict that the Neanderthal-derived allele weakens CTCF binding leading to reduced insulation between *ICOS*, a T-cell costimulator, with downstream contacts.

### 2.4 Regions with 3D divergence highlight AH-MH phenotypic differences

To explore the functional effects of AH-MH 3D genome divergence, we tested for phenotypic annotation enrichment. We considered the 43 loci with shared divergence between MHs and all three Neanderthals (Fig. 3B). Although the loci were identified at approximately 1 Mb resolution, most 3D modifications disrupt a smaller sub-window. Thus, as described in the example above (Fig. 3C), we used *in silico* mutagenesis to identify the AH-MH sequence change(s) that produced the largest disruption in the contact maps. We will refer to these as “3D-modifying variants” (Methods). We then intersected the predicted 3D-modifying variants with experimentally defined TADs to determine the genes to which they are physically linked. Ultimately, we found 88 physical links to protein-coding genes (85 unique genes) for the 45 3D-modifying variants in the 43 Neanderthal-MH 3D divergent loci (Tables S2,S5).

We tested if these genes are enriched for phenotypic annotations using both gene-phenotype links from rare disease (OMIM Human Phenotype Ontology [HPO] terms) and common disease databases (GWAS Catalog 2019) [89–93]. 3D genome organization perturbation has been linked to both types of disease: large-scale disruption leading to severe disease and subtle changes in regulatory insulation contributing to complex traits disease [69–72, 74]. We find links to 271 and 208 candidate traits from the rare and common disease ontologies, respectively. For each trait, we test if the observed overlap with 3D divergent loci is more than expected by chance using an empirically-generated null distribution (Methods). In summary, this sequential process links 3D divergent windows to variants to TADs to genes and, ultimately, phenotypes (Fig. S8).

With the HPO annotations, we found enrichment for effects of these genes related to the eye (retinopathies, optic atrophy, constricted visual field [most significant association: 27× enriched, *P* = 2 × 10^−5^]), skeletal system (notably, supraorbital ridge morphology [12×, *P* = 0.002]), and hair (e.g. low anterior hairline [12×, *P* = 0.003]) (Fig. 3D, top). In the GWAS Catalog annotations, we find enrichment related to intelligence and cognition (13×, *P* = 0.0002), lung function (NO levels, COPD [35×, *P* = 0.0008]), response to certain medications (30×, *P* = 0.002), immunologic response (ankylosing spondylitis, allergy, inflammatory bowel disease [12×, *P* = 0.004]), and brain region volumes (putamen, subcortex [17×, *P* = 0.006]) (Fig. 3D, bottom). Trait enrichments for 3D-modifying variants found in Denisova are highlighted in Fig. S9. Because Denisova and Neanderthal share many alleles, some similar traits are enriched (retinopathy, intelligence, lung function, etc.); however, overall, we find fewer enriched traits.

In summary, genomic loci with 3D divergence between Neanderthals and MHs are enriched for physical proximity to genes associated with a diversity of traits related to the skeleton, eye, hair, lung, immune response, brain region volume, and cognitive ability. These findings align with and expand what we know from both the fossil-record and previous work based on variants in MHs [11, 14–20]. Importantly, our approach permitted the interrogation of variants unobserved in MHs (76% of predicted 3D-modifying variants), and it provides a putative molecular mechanism for the phenotypic differences.

### 2.5 Relationship between sequence divergence and 3D divergence

Given that we observe 3D differences between AH and MH genomes, we quantified the relationship between 3D and sequence divergence on both genome-wide and more local scales. First, we computed the genome-wide 3D genome divergence for all pairs of AH and MH individuals. We find the mean 3D genome divergence largely follows sequence divergence (Figs. 4A,S10). Neanderthals are the most similar in 3D genome organization to other Neanderthals, then to the Denisova, and then to MHs (mean 3D divergences: 9.8×10^−4^, 3.4×10^−3^, and 4.3×10^−3^, respectively). Genome-wide 3D divergence also tracks with sequence divergence within the Neanderthal: Vindija and Chagyrskaya are more similar than they are to the outgroup Altai (Vindija-Chagyrskaya mean 3D divergence of 8.4 × 10^−4^ vs. Vindija-Altai of 1.0 × 10^−3^) [3].

**Figure 4:**
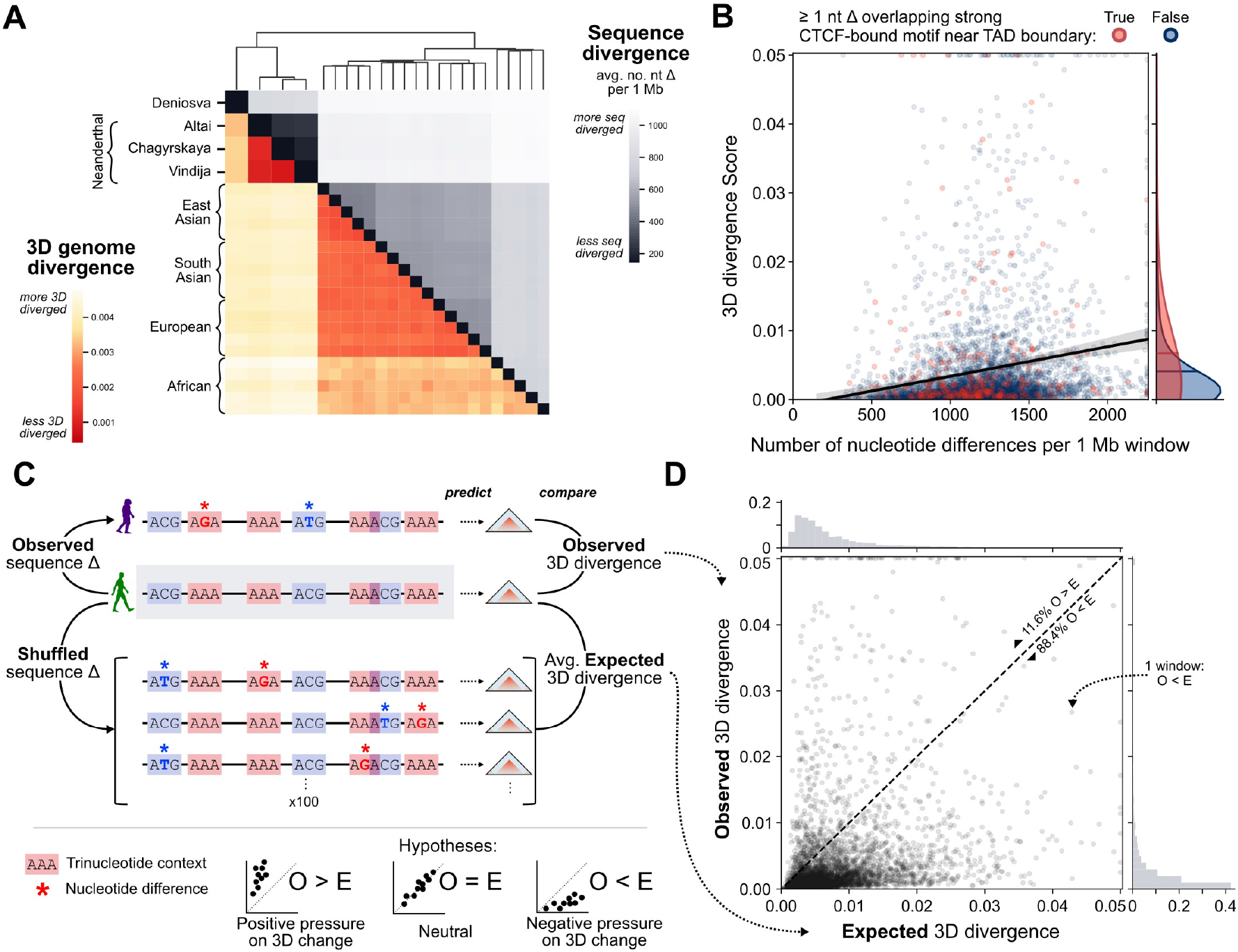
3D genome organization constrained human sequence divergence. **(A)** 3D genome divergence (lower triangle) follows patterns of sequence divergence (upper triangle). AHs have more similar 3D genome organization to each other than to 15 MHs from different 1000G super-populations. Clustering is based on sequence divergence; see Fig. S10 for clustering by 3D genome divergence and data for each sub-population. **(B)** Sequence divergence is only very modestly correlated with 3D genome divergence (*r*^2^ = 0.011, *P* = 2.3 × 10^−13^, *N* = 4999). Each point represents a 1 Mb window from a genome-wide comparison between the 3D genome organization of a Neanderthal (Vindija) and African MH (HG03105) individual and the black line with band represents a linear regression with 95% CI. Windows with large 3D divergence are enriched for MH-AH nucleotide (nt) differences overlapping a strong CTCF-bound motif within 15 kb of a TAD boundary (red) (two-tailed Mann–Whitney U *P* = 0.00077). **(C)** To evaluate whether 3D genome organization constrained sequence divergence, we estimate the null distribution of expected 3D divergence based on sequence differences between the Neanderthal (Vindija) and African MH (HG03105) genomes. We shuffle observed nucleotide differences (stars) while preserving tri-nucleotide context (colored rectangles) and predict 3D genome organization for 100 shuffled sequences for each window. Under a model of no sequence constraint due to 3D organization, observed 3D divergence would equal the expected 3D divergence (*O* = *E*). Alternatively, observing more 3D divergence than expected would suggest positive selection on sequence changes that cause 3D divergence (*O* > *E*). Finally, observing less 3D divergence than expected would suggest negative pressure on sequence changes that cause 3D divergence (*O < E*). **(D)** Observed 3D divergence is significantly less than the mean expected 3D divergence based on sequence (*O* < *E*: 88.4% of *N* = 4, 999 windows below the diagonal, binomial-test *P <* 5 × 10^−324^). The mean expected 3D divergence is on average 1.78-times higher than the observed 3D divergence (*t*-test *P* = 1.8 × 10^−48^). 3D divergence scores greater than 0.05 and nucleotide differences greater than 2250 are clipped to the baseline for visualization purposes.

Next, we evaluated if sequence divergence and 3D divergence are correlated on the local scale. We find a very weak positive relationship between 3D and sequence divergence at the 1 Mb window level (Fig. 4B, *r*^2^ = 0.01, *P* = 2.3 × 10^−13^). As suggested by the weak correlation, many windows with low sequence divergence have high 3D divergence, and many windows with high sequence divergence have low 3D divergence.

Given the weak relationship between sequence and 3D divergence, we sought to identify some properties of sequence differences that result in large 3D divergence. Based on the importance of CTCF-binding in maintaining 3D genome organization [50, 62, 79, 80], we quantified the effects of AH-MH nucleotide differences overlapping CTCF binding motifs. Disruption of CTCF binding sites is important, but not all disruptions are likely to influence 3D divergence. Leveraging additional functional genomics data on CTCF binding and TAD boundaries, we find that the quantity, quality, and context (e.g., strength of a motif and proximity to a TAD boundary) influence whether AH-MH sequence divergence will result in a 3D organization divergence (Fig. S11). For example, if a window has at least one AH-MH nucleotide difference overlapping a strong CTCF-bound motif near a TAD boundary (within 15 kb), the AH-MH 3D divergence is 1.64-times greater (*P* = 0.00077, *N* = 260*/*4999 windows, Fig. 4B). Thus, we are observing complex sequence patterns underlying 3D genome folding that could not be determined by simply considering sequence divergence or intersecting AH variants with all CTCF sites. This is concordant with previous results which suggest that 3D genome folding is governed by a complex CTCF binding grammar [50, 80, 82, 83].

### 2.6 Maintenance of 3D genome organization constrained sequence divergence in recent hominin evolution

Next, we evaluated if the pressure to maintain 3D genome organization constrained recent human sequence evolution. We estimated whether the amount of 3D divergence between AHs and MHs is more or less than expected given the observed sequence divergence. To compute the expected 3D divergence distribution for each 1 Mb window, we shuffled observed nucleotide differences between an African MH (HG03105) and AH (Vindija Neanderthal) 100 times and applied Akita to predict the resulting 3D genome divergence (Fig. 4C). We controlled for the non-uniform probability of mutation across sites using a model that preserved the tri-nucleotide context of all variants in each window with each shuffle. For each 1 Mb window, we compared the observed 3D divergence with the expected 3D divergence from the 100 shuffled sequences with the same nucleotide divergence.

If the 3D genome does not influence sequence divergence, the observed 3D divergence would be similar to the expected 3D divergence (Fig. 4C, bottom-middle). Alternatively, if the observed 3D divergence is greater than expected based on sequence divergence (Fig. 4C, bottom-left), this suggests positive selection on variation contributing to 3D differences. Finally, if the observed 3D divergence is less than expected based on sequence divergence (Fig. 4C, bottom-right), this suggests negative pressure on variation contributing to 3D differences.

We find that observed 3D divergence is significantly less than expected based on sequence divergence (Fig. 4D). 88.4% of 1 Mb windows have less 3D divergence that expected based on their observed sequence differences (binomial-test *P* < 5 × 10^−324^). Genome-wide, the mean expected 3D divergence is 78% higher than the observed 3D divergence (*t*-test *P* = 1.8 × 10^−48^). This suggests that, in recent hominin evolution, pressure to maintain 3D genome organization constrained sequence divergence. This aligns with previous studies that demonstrated depletion of variation at 3D genome-defining elements (e.g., TAD boundaries, CTCF sites) [73–77], but it specifically implicates 3D genome folding.

### 2.7 3D genome organization constrained introgression in MHs

Eurasian individuals have on average 2% AH ancestry due to introgression; however, AH ancestry is not evenly distributed throughout the genome [2, 15, 31]. Our previous analyses demonstrate that AH and MH exhibit a range of 3D genome organization divergence across the genome (Fig. 2C) and that pressure to maintain 3D genome organization constrained sequence divergence (Fig. 4D). Thus, we hypothesized that for a given genomic window, its tolerance to 3D genome organization variation in MHs would influence the probability that introgressed AH DNA is maintained in MH.

To test this, we first quantified the levels of 3D genome diversity for 20 modern Africans in 1 Mb sliding windows across the genome. We then computed the average African-African 3D genome divergence and term this “3D genome variability”. Genomic windows with low 3D genome variability have similar 3D genome organization among all Africans, suggesting these loci are less tolerant of 3D folding changes. In contrast, regions with high 3D genome variability suggest a diversity of 3D genome organization present. Finally, we computed the amount of introgressed sequence in Eurasian populations for each window (Methods, [94]).

Genomic windows with high levels of introgression across Eurasians are enriched for windows with higher 3D genome variability (Fig. 5A, Mann-Whitney U *P* = 0.0007). On average, windows with evidence of introgression have 72% higher 3D genome variability than windows without introgression. Moreover, the magnitude of 3D genome variability is predictive of the average amount (proportion of bp) of introgressed sequence remaining in a 1 Mb window (*P* = 5.7 × 10^−9^, Fig 5B, vertical axis). As expected, sequence variability is very significantly predictive of the amount of introgressed sequence in a window (*P* = 1.9 × 10^−49^, Fig 5B, horizontal axis). Yet, even when conditioning on sequence variability, 3D genome variability provides additional information about the amount of AH ancestry in a window (Fig 5B, conditional *P* = 5.7×10^−4^). In other words, even if two windows have the same level of sequence variability in MHs, windows that are more 3D variable are more likely to retain introgressed sequence. We also find that 3D genome variability is more strongly predictive of introgression shared among all three super-populations than an introgressed sequence unique to a single super-population (Supplemental Text, Tables S7,S8). Using earlier introgressed Neanderthal haplotype predictions from Vernot et al. [15] and other thresholds yield similar results (Figs. S12,S13). Because we compute variability in Africans with very low levels of AH ancestry, the increased 3D genome variability in MHs is not a result of introgression.

**Figure 5:**
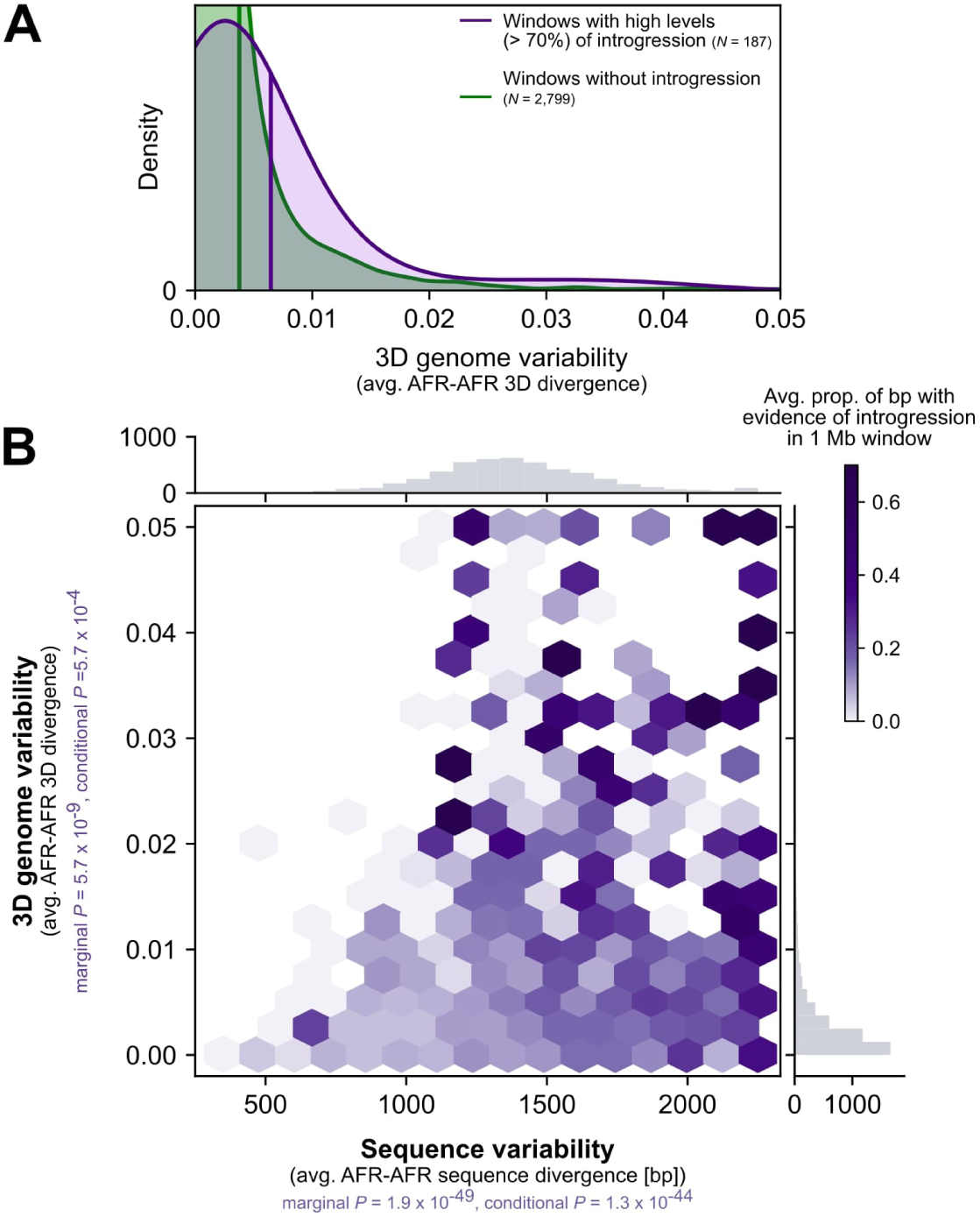
3D variable windows in MH have more evidence of AH introgression. **(A)** Windows with high levels of introgression across present-day non-African populations (purple, *N* = 187) are more 3D-variable in modern Africans (horizontal axis) than windows without evidence of introgression (green, *N* = 2, 799; two-tailed Mann–Whitney U *P* = 0.0007). Vertical lines represent the distribution means. Introgression is called based on Sprime [94]. To focus on regions consistently tolerant of AH ancestry, we considered introgression shared across 1000 Genomes super-populations and covering at least 70% of bases in a 1 Mb window (Methods). Results from other introgression sets and thresholds are similar (Figs. S12–S13 and Tables S7–S8). **(B)** The relationship between sequence variability (horizontal axis) and 3D genome variability (vertical axis) with amount of AH ancestry in a window. Darker purple indicates a higher proportion of introgression in a 1 Mb genomic window. Sequence variability (*P* = 1.9 × 10^−49^) and 3D genome variability (*P* = 5.7 × 10^−9^) both independently predict amount of introgression. Additionally, even when controlling for sequence variability in a window, 3D genome variability is informative about the amount of introgression (*P* = 5.7 × 10^−4^).

These results suggest that 3D genome organization shaped the landscape of AH introgression in modern Eurasian genomes. Previous findings demonstrated Neanderthal ancestry is depleted in regions of the genome with strong background selection, evolutionary conservation, and annotated molecular function (e.g. genes and regulatory elements) [11, 30, 31, 40, 41]. Our results expand this to implicate the 3D genome as a contributor to the landscape of AH ancestry in MHs today.

### 2.8 Introgression shaped the 3D genome organization of present-day Eurasians

Given the differences between AH and MH 3D genome organization at many loci, we hypothesized that introgressed AH sequences could have introduced novel 3D contact patterns to Eurasian MHs. To test this, we integrated Eurasians into our previous comparisons of AHs and African MHs.

For example, we found an AH-MH 3D divergent window on chromosome 7 with a striking pattern of 3D genome diversity across modern Eurasians (Fig. S14). The 3D genome divergence between all Africans and AH was consistently high, as required to be an AH-MH divergent locus. Out of 15 representative Eurasians, 11 had divergent organization compared to the Neanderthal 3D contact map. However, four Eurasians had very low 3D divergence from the Neanderthal indicating similar 3D organization. We hypothesized that these Eurasians inherited Neanderthal ancestry at this loci via introgression, and that the Neanderthal alleles produced the novel pattern of 3D genome organization.

When examining the contact maps of this window, all Africans have a large approximately 450 kb loop domain starting near the promoter of *IGFBP3*, a gene encoding insulin-like growth factor binding protein 3 (Fig. 6A). In contrast, Neanderthals (Vindija, Chagyrskaya, and Altai) have two smaller sub-domains insulated by a CTCF site. Using *in silico* mutagenesis, we identify that the variant with the largest effect on 3D organization is a G to A change at chr7:46,169,621 (rs12536129). The derived A allele, which strengthens the CTCF motif, appeared along the Neanderthal lineage. The Eurasians with 3D genome organization very similar to Neanderthals all have an introgressed haplotype carrying the Neanderthal-derived A allele overlapping this CTCF site, whereas individuals without introgression harbor the predicted ancestral genome organization (Fig. S14)[96]. In summary, we identified a putative novel 3D genome pattern inherited by modern Eurasians from Neanderthals at this loci.

**Figure 6:**
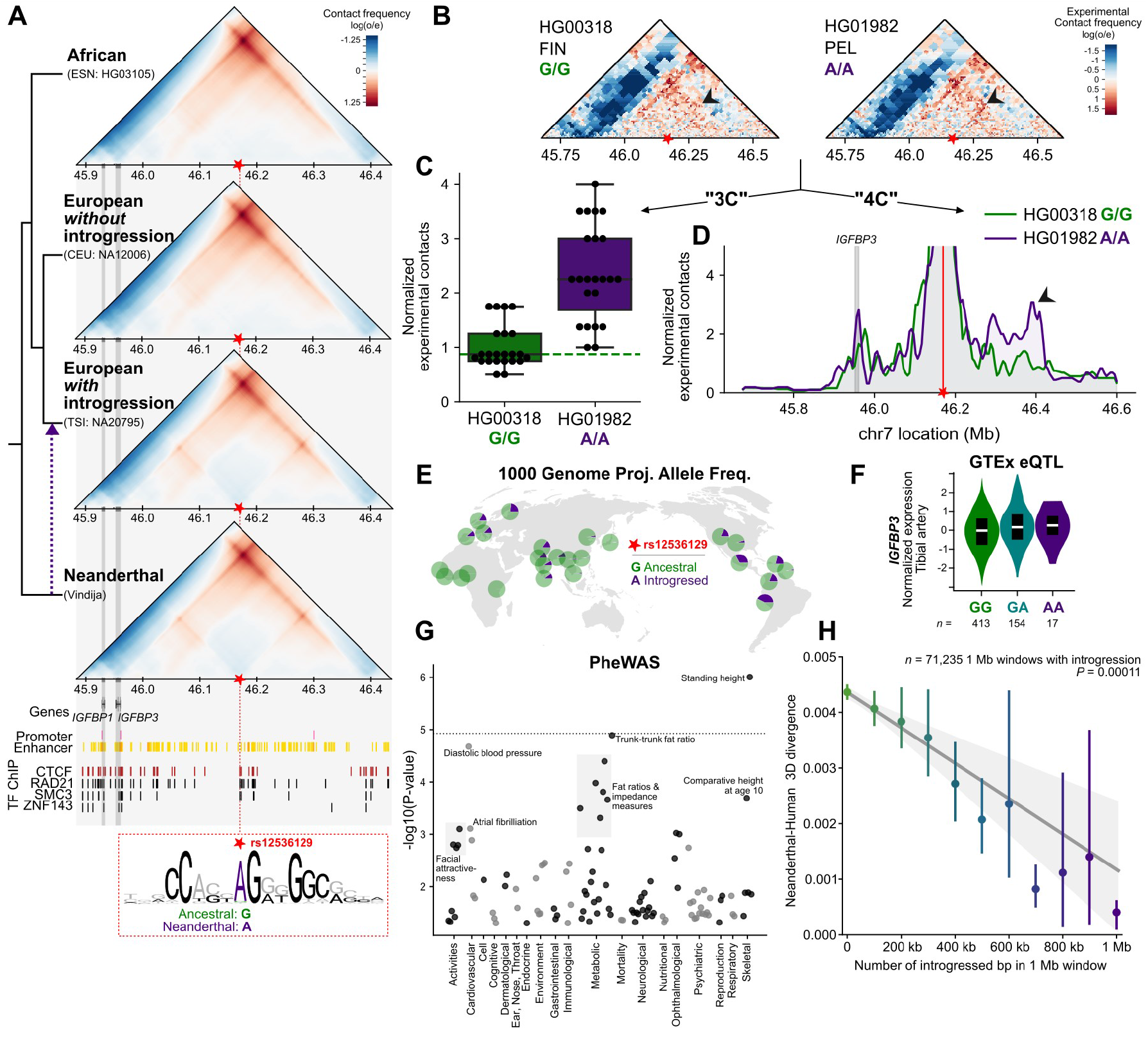
Introgression introduced novel 3D genome organization patterns to modern Eurasians. **(A)** Locus on chr7 (chr7:45,883,392-46,436,352) where Eurasians with introgression inherit 3D genome organization patterns from Neanderthals. Neanderthals and individuals with introgression have two domains insulated by a CTCF site (red box). In MHs without introgression, this motif is disrupted with a G instead of an A (star, chr7:46,169,621, rs12536129) leading to a larger fused domain. **(B)** Hi-C experimental validation of this locus in an individual homozygous for the ancestral allele (HG00318, G/G) and homozygous for the introgressed allele (HG01982, A/A) recapitulates the computationally-predicted substructure. **(C)** The subset of contacts predicted to be most divergent using the computational maps demonstrate significantly increased 3D contact in the experimental Hi-C maps (*P* = 2.0 × 10^−7^, paired T-test). **(D)** The individual with introgression (HG01982) demonstrates greater experimental contacts downstream to the CTCF site (black arrowhead) and at the location of *IGFBP3* concordant with the predicted increased substructure. **(E)** Across modern Eurasian populations in the 1000 genome project, this introgressed allele is at high-frequency. **(F)** This introgressed allele is an eQTL in GTEx for the physically linked gene *IGFBP3* (*P* = 0.00014 in artery tissue) [42]. **(G)** In MHs, this variant is associated with traits including standing height (*P* = 9.9 × 10^−7^), fat distribution (trunk fat ratio, impedance measures, *P* = 1.3 × 10^−5^), and diastolic blood pressure (*P* = 2.1 × 10^−5^) [95] (dotted line is a highly conservative Bonferroni-corrected *P* -value). **(H)** The amount of introgression in a 1 Mb window (number of bp, horizontal axis) is significantly correlated with the similarity of an individual’s 3D genome organization to a Neanderthal’s (Vindija) genome organization (vertical axis) (*P* = 0.00011, *n* = 71, 235 1 Mb windows across 15 Eurasians). The error bars signify 95% bootstrapped CIs and the error band signifies the 95% bootstrapped CI for the linear regression estimate.

To validate this putative Neanderthal-derived 3D organizational pattern experimentally, we identified individuals in the 1000 Genomes Project with each possible genotype at this locus and performed Hi-C on their LCLs. We analyzed: a Finnish individual (HG00318) homozygous for the ancestral allele (G/G); a Peruvian individual (HG01982) homozygous for the introgressed allele (A/A); and an individual from Utah (GM12763) heterozygous (G/A) at this locus (Fig. S15). Consistent with the 3D genome predictions, the individual with introgression demonstrates the novel Neanderthal-like 3D genome substructure in their Hi-C map (Fig. 6B).

To quantify these differences in genome structure, we tested a subset of Hi-C contacts within the “all-versus-all” 3D map using both a “one-versus-one” strategy akin to 3-C and an “all-versus-many” strategy akin to 4-C. In the “one-versus-one” strategy, we identified the subset of contacts which were *a priori* predicted to be most divergent using the computationally determined maps (Methods, Fig. S15B-C). We confirm that this subset of contacts in the experimental Hi-C maps is significantly stronger in the individual with introgression than the individual without introgression at this locus (*P* = 2.0 × 10^−7^, Fig. 6C). In the “one-versus-many” strategy, we quantified contacts between the location of the modified CTCF motif and the rest of the locus. The individual with introgression demonstrates greater experimental contacts downstream to the CTCF site concordant with the predicted increased substructure. They also harbor a stronger peak of contact between the CTCF site and *IGFBP3*, Fig. 6D). The individual heterozygous for Neanderthal ancestry at this locus has a pattern of 3D contact in between the homozygotes (i.e. substructure stronger than the homozygous ancestral but weaker than the homozygous introgressed genotype, Fig. S15).

After demonstrating the 3D changes associated with the introgressed allele, we sought to examine the possible functional consequences for gene regulation and phenotypic trait variation. This introgressed allele is common in non-African populations, but it varies substantially in frequency across populations (28% AMR, 2% EAS, 16% EUR, 11% SAS, 0% AFR), with highest frequency in Peru at 42% (Fig. 6E). We find that this variant is an eQTL for the gene *IGFBP3*, which has increased physical contact with the CTCF site (Fig. 6F, *P* = 0.00014 in GTEx artery tissue) [42]. Underscoring the importance of the Neanderthal allele in MHs, this variant is associated with traits including standing height (*P* = 9.9 × 10^−7^), fat distribution (trunk fat ratio, impedance measures, *P* = 1.3 × 10^−5^), and diastolic blood pressure (*P* = 2.1 × 10^−5^) (Fig. 6G).

Given these examples of Neanderthal introgression contributing novel 3D folding to present-day Eurasians, we searched for similar patterns genome-wide. We considered 4,749 autosomal 1 Mb windows for 15 Eurasians (total *n* = 71, 235) to quantify the relationship between the amount of introgression and 3D similarity to Neanderthals. We find that the amount of introgression (bp per window) is significantly correlated with 3D divergence to the Vindija Neanderthal (*P* = 0.00011, Fig. 6H). Results from comparisons to the other Neanderthals are consistent (Fig. S16). On average, in a 1 Mb window, if an individual has 80% Neanderthal ancestry, their 3D genome is 2.4 times more similar to the Neanderthal 3D genome than if they have no (0%) Neanderthal ancestry.

In summary, Eurasians with more Neanderthal ancestry in a window have more Neanderthal-like 3D genome folding patterns. Furthermore, at an example locus, we identify a novel 3D organization pattern inherited by modern humans and validate this using Hi-C, highlighting a putative molecular mechanism for the effect of Neanderthal ancestry on human gene expression and trait variation.

## 3 Discussion

The role of 3D genome organization in human biology is increasingly recognized [62, 73–77]; however, current techniques for measuring 3D folding cannot be applied to the study of ancient DNA. Furthermore, despite methodological improvements in assays of the 3D genome, high-resolution experiments across many diverse individuals, species, and cell types remain cost prohibitive [97, 98]. To address these gaps, we provide a framework for inferring 3D genome organization at population-scale in ancient and modern genomes that facilitates evaluation of previously untestable hypotheses.

First, we apply this framework to resurrect archaic 3D genome organization. We find that 3D genome organization constrained sequence divergence and patterns of introgression in hominin evolution. We catalog genomic regions where AH and MH 3D genome organization diverged and illustrate how this novel mechanism links sequence differences to phenotypic differences. Importantly, our approach permitted the evaluation of variants unobserved in MHs, and it provides a putative molecular mechanism for AH-MH phenotypic differences including those that may have been selected against after hybridization (e.g. cognitive and brain morphology traits) [11, 19, 30, 31, 39–41]. Finally, we identify regions in which introgression introduced archaic 3D genome folding that are novel to Eurasians and then use Hi-C to validate 3D genome divergence in a high-frequency introgressed region. Together, these results illustrate the power of imputing unobservable molecular phenotypes to resolve evolutionary questions about functional divergence.

Second, we anticipate that our framework for comparing and interpreting hundreds of genome-wide 3D genome contact maps will be helpful for testing hypotheses beyond archaic DNA. In the interpretation of genetic variants of unknown significance, it will be key to consider the effect of inter-individual and interspecies variation on 3D genome architecture, especially given recent evidence that even common DNA sequence variants can influence 3D organization and human phenotypic variation [72, 97–99]. Our work establishes the groundwork to answer many diverse questions. For example, we illustrate how *in silico* mutagenesis can highlight the role of a variant in 3D genome organization and how to integrate this with other functional annotations. This allows us to examine the 3D effects of variants never before observed in MHs, which is essential to non-coding variant interpretation from the lens of both evolution and disease. Our measure of “3D genome variability” provides genome-wide quantification of how different regions tolerate variation in 3D genome folding, and it could be integrated with existing metrics for quantifying differences in 3D contact maps [88]. We also demonstrate a simulation approach for testing how 3D genome folding constrains sequence evolution across the genome. Finally, we develop a method to robustly identify 3D divergent windows between populations. With the recent growth of 3D genome *in silico* predictors [81–84, 100, 101], we anticipate that our work can provide a foundation for both hypothesis generation and prioritization of experimental resources. Recent work has demonstrated use of this framework for *in silico* investigation of the role of repetitive elements in 3D genome organization [102], in quantifying the role of alternative splicing in the divergence of archaic hominins [99], and for exploring the diversity of 3D genome folding across diverse populations of MHs [97] and non-human primates [98]. Although our approach provides many novel benefits, it also has limitations that we hope future work will address. First, our comparisons likely underestimate 3D diversity. We only investigate windows of the genome with complete sequence coverage. Because of ancient sample degradation, we do not have full coverage of AH genomes. We use a conservative approach to effectively mask regions of the genome lacking coverage in AHs (Fig. S1 and Methods). Furthermore, we only consider the effects single nucleotide variants. We do not consider structural variation (SV) due to the challenges of calling SV accurately in ancient samples. We anticipate new methods in ancient DNA sequencing will allow us to model the 3D genome organization of AHs more completely. Second, our 3D genome organization comparisons are based on a simple correlation-based metric. Correlation-based metrics are a reliable first step for identifying divergence [88] and we demonstrate concordance using other more biologically informed methods (Fig. S4). Third, although Akita is trained simultaneously across five cell types, 3D genome organization is largely conserved across cell types and predictors only identify limited cell-type-specific differences. Therefore, we focused on the highest resolution predictions in a single context (HFF). As more high-resolution Hi-C and Micro-C become available across diverse cell types, our framework can be applied to identify cell-type-specific AH-MH differences.

Several practical caveats must be considered when interpreting some of our results. For example, to conduct *in silico* mutagenesis we manipulate every single nucleotide separately against the same background rather than considering the prohibitively large number of possible variant combinations. Additionally, we prioritize the investigation of *cis*-regulatory variation rather than long-range combinatorial variation that likely contributes to the robustness of gene regulation [103]. The annotations that link 3D-modifying variants to genes and functions are also based on studies in MHs (HPO and GWAS). It is possible, though unlikely, that a gene disrupted in MHs would lead to different traits in AHs. Finally, while we generate new Hi-C data for three individuals with differential introgression at a locus of interest, given the scope of our study and the nature of archaic DNA, direct experimental validation at scale is not feasible. However, we use complementary experimental data (CTCF ChIP-seq, experimentally-derived TADs, eQTL) to provide independent support for the influence of variants on 3D genome organization and to link variants with genes in true physical proximity. Even if high-resolution Hi-C were available across many Eurasians, an experimental approach would still not capture all AH variation, highlighting the necessity of our computational approach.

In conclusion, our framework for inferring archaic 3D genome organization provides a window into previously unobservable molecular mechanisms that shaped the sequence and phenotypic evolution of hominins.

## 4 Methods

### 4.1 Modern human and archaic genomes

#### Obtaining genomes

All genomic analysis was conducted using the GRCh37 (hg19) genome assembly and coordinates (www.ncbi.nlm.nih.gov/assembly/GCF_000001405.13/). Genomic variation within modern humans (MH) came from 1KGP, Phase 3 from Auton et al. [87]. All MH genomes were selected randomly from each subpopulation with a filter for females only to facilitate comparisons of the X chromosome. The 1000 Genomes Project (1KGP) individuals used are listed in Table S1. Archaic genomes are from Prüfer et al. [1] (Altai), Prüfer et al. [2] (Vindija), Mafessoni et al. [3] (Chagyrskaya), and Meyer et al. [4] (Denisova).

#### Building individual genomes

We constructed full-length genomes for each MH or AH based upon the genotyping information in their respective vcf file. Given the difficulty of distinguishing heterozygous genotypes in the ancient DNA samples, we treated all individuals as if they were homozygous (pseudo-haploid). We built each individual genome using GATK’s FastaAlternateReferenceMaker tool [104]. If an individual had an alternate allele (homozygous or heterozygous), we inserted it into the reference genome to create a pseudo-haploid, or “flattened” genome for each individual. This procedure is illustrated in step 1 of Fig. S1.

#### Accounting for missing data in the archaic genomes

Ancient DNA is both fragmented and degraded. These characteristics present challenges to both sequencing and alignment, resulting in gaps in coverage, particularly in genomic regions of low complexity. To account for this missing data, we “masked” all genomic regions lacking archaic genotyping information by reverting nucleotide states to the hg19 reference. For analyses that compared 3D genome organization between MHs and AHs, we masked both MH and AH genomes. This procedure is illustrated in steps 2-4 of Fig. S1. Archaic genome coverage is shown in Fig. S2. For analyses that only considered MHs (e.g. quantifying 3D genome variability across the genome in MHs), this masking procedure was not applied.

### 4.2 3D genome organization predictions with Akita

After the genomes were prepared, we input them into Akita for predictions using a 1 Mb sliding window (1,048,576 bp) overlapping by half (e.g. 524,288-1,572,864, 1,048,576-2,097,152, 1,572,864-2,621,440). Although Akita is trained simultaneously on Hi-C and Micro-C across five cell types in a multi-task framework to achieve greater accuracy, we focus on predictions in the highest resolution maps, human foreskin fibroblast (HFF). We note that the results are similar when considering other cell types (e.g. embryonic stem cells), likely because of limited cell-type-specific differences (Fig. S5). Akita considers the full window to generate predictions, but the resulting predictions are generated for only the middle 917,504 bp. Each contact map is a prediction for a single individual, and each cell represents physical 3D contacts at approximately 2 kb (2,048 bp) resolution. The value in each cell is log_2_(*obs/exp*)-scaled to account for the distance-dependent nature of chromatin contacts. Darker red pixels indicate more physical contacts and darker blue pixels denote fewer physical contacts. For all analyses, we only considered windows with full (100%) coverage in the hg19 reference genome for a total of 4749 autosomal and 250 chromosome X windows. Fudenberg et al. [82] provides further details on the CNN architecture and training data used.

### 4.3 3D genome comparisons

After predictions were made on all 1 Mb windows for all individuals, we compared the resulting predictions using a variety of measures. All measures are scaled to indicate divergence: higher indicates more difference while lower indicates more similarity. In the maintext we transform the Spearman’s rank correlation coefficient (1 − *ρ*) to describe 3D divergence. We consider measures based on the Pearson correlation coefficient (1−*r*) and mean squared difference 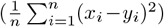 in Fig. S4. Percentiles of 3D divergence shown in Fig. 2A-B are calculated with reference to a universe of 4 AHs × 5 African MHs × 4999 genomic windows for a total of 99,980 comparisons. Figs. 4A,S10 averages the 3D divergence (1 − *ρ*) across all 4999 1 Mb windows (lower triangle) to compare to the average number of bp differences (after the masking procedure described above) in the same pair of individuals (upper triangle). Clustering is done with the “complete” (Farthest Point) method.

### 4.4 AH-MH 3D divergent loci

#### Identifying loci

To identify loci with AH-MH 3D genome organization divergence, we compared the 3D contact map at each 1 Mb loci between each AH and 20 African MHs. To call a region as divergent, we required all 20 AH-MH comparisons to be more 3D divergent than all MH-MH comparisons (Fig. 3A). This identifies regions with consistent 3D differences between AHs and MHs while excluding regions with a large 3D diversity in modern humans. We also required the minimum AH-MH 3D divergence to be in the 80^th^ percentile or greater of most 3D diverged (Fig, 2A, 3D divergence > 0.0042). Because 20 MHs do not capture the full MH genome diversity, it is possible that these criteria would still capture 3D patterns segregating in modern Africans that are not truly AH-MH diverged. Thus, we removed any windows where the 3D-modifying variant determined by *in silico* mutagenesis (below) was observed in 1KGP MHs if it was not introgressed (LD of *r*^2^ = 1 with introgressed variants called by Browning et al. [94] or Vernot et al. [15]). For the counts of AH-MH divergent windows (Fig. 3B), we considered overlapping 1 Mb windows as a single observation. We summarize and report the AH-MH 3D divergent windows in Tables S2,S3 and a larger set of windows based on less conservative criteria in Table S4.

#### *In silico* mutagenesis

To identify the variant(s) contributing to the most prominent 3D differences in each identified AH-MH divergent window, *3D-modifying variants*, we use *in silico* mutagenesis. For example, for an Altai Neanderthal divergent window, we identify every bp difference that is unique to the Altai genome when compared to 20 African MH genomes. In the background of the MH (HG03105) genome, we insert each different Altai allele one-at-a-time. We then compare the resulting contact map between the original MH genome and the MH genome with each Altai allele. We then identify both the allele resulting in the largest 3D divergence and any other variants that contribute to a 3D divergence >= 0.0042 and term these “3D-modifying variants” (Table S2,S5).

#### Phenotype ontology enrichment

To test if AH-MH 3D-modifying variants are enriched near genes related to particular phenotypes we follow a procedure visually described in Fig. S8. 3D-modifying variants (above) are linked to genes in their TAD because this provides evidence of physical proximity. TADs are defined as regions between TAD boundaries as defined with MicroC data in HFF from Akgol Oksuz et al. [105] (lifted over to hg19). Genes are defined as the longest transcript from protein-coding genes (NM prefix) from NCBI RefSeq downloaded from the UCSC Table Browser [106]. Genes are linked to phenotypes from the Human Phenotype Ontology (HPO) and GWAS Catalog 2019 downloaded from Enrichr [91–93]. Annotations are further grouped into phenotypic systems via system-level annotations from Gene ORGANizer [107] and manual curation. HPO largely considers rare disease annotations and has 1779 terms with 3,096 genes annotated [89]. The GWAS Catalog largely considers common disease annotations and has 737 terms with 19,378 genes annotated [90]. Through this procedure, we counted the number of ontology terms linked to the set of 3D-modifying variants. We considered 3 different sets, those shared (intersect) by all Neanderthals (Fig. 3), those in any Neanderthal (union), and those in Denisova (Fig. S9, Table S2).

We test enrichment for ontology terms linked to at least one 3D-modifying variant. While the annotations are downloaded from Enrichr, we did custom enrichment analyses with a more appropriate null. For each set, we shuffle the TADs that harbor each 3D-modifying variant into the background genome. We defined the background genome as any place where a 3D-modifying variant could have been identified (i.e. regions with full coverage in modern humans used for Akita predictions). We then use the same procedure (Fig. S8 to link the shuffled TADs to genes and then ontology terms. We repeat this shuffle 500,000 times to create an empirical distribution for how many times we would observe each annotation linked to a 3D-modifying variant under the null. We used these distributions to calculate an enrichment and *P* -value for each ontology term. The FDR-corrected significance level was determined empirically using these null observations (a subset of *n* = 50, 000). We select the highest p-value threshold that led to a *V/R* < *Q* where *V* is the mean number of expected false discoveries and *R* is the observed discoveries (which includes both true and false positives).

### 4.5 Sequence comparisons

We compare 3D genome divergence with sequence divergence in Figs. 4. To calculate the sequence divergence between two individuals, we counted the proportion of bases at which the two individuals differ in the 1 Mb window. For comparisons of divergence when including AHs, we applied the same masking procedure as used to facilitate 3D genome comparisons (i.e. windows with missingness in AHs are filled with hg19 reference).

### 4.6 CTCF motif overlap

We consider how nucleotide differences in a window (between Neanderthal [Vindija] and an African MH [HG03105]) impacts 3D genome divergence in Figs. 4B,S11. We stratified variants by if they overlap a bound CTCF motif and their distance to TAD boundaries. CTCF motifs are from Vierstra et al. [108]. CTCF-bound open chromatin candidate cis-regulatory elements (cCREs) in the HFF cell type are from Abascal et al. [109]. TAD boundaries in the HFF cell type are from processed MicroC data from Akgol Oksuz et al. [105]. These annotations were all lifted over to hg19 [110]. A window was considered to have a CTCF-overlapping variant if an AH-MH nucleotide difference intersected a CTCF-bound HFF cCRE and a CTCF motif. Results were further stratified by varying levels of motif strength (“match score” in the top 10^th^,25^th^, 50^th^, or any percentile), distance to TAD boundary (within 15 kb, 30 kb, or anywhere), and whether the CTCF motif overlap occurs in the middle 50% of the 1 Mb window or not.

### 4.7 Empirical distribution of expected 3D genome divergence

To compute the expected 3D divergence in a window given the observed sequence divergence, we generate genomes with shuffled nucleotide differences. We match these shuffled differences to the same number and tri-nucleotide context of the observed sequence differences between the Neanderthal (Vindija) and an African MH (HG03105) genome (Fig. 4C). Variants are not shuffled into masked regions of the genome. For each 1 Mb window of the genome (*N* = 4999) we generate 100 shuffled sequences. We calculate an empirical distribution of expected 3D divergence from comparing the contact maps of the shuffled sequences with the MH sequence. Finally, we compare the average expected 3D divergence from this distribution to the observed AH-MH 3D divergence.

### 4.8 Relationship between 3D genome organization and introgression

#### 3D genome variability

To consider how 3D organization may have constrained where we observe introgression in the genome, we calculated 3D genome variability across the genome in MHs. Because we are not comparing these predictions with AH 3D genome organization, we did not mask the genomes before 3D genome predictions (above). In the same 1 Mb sliding windows across the genome, we predicted the contact maps for 20 modern Africans (because they have no or very little introgression). For each window, we calculate the 3D genome divergence between all 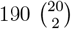 pairs of contact maps. We then computed the “3D genome variability” by taking the mean of these 190 divergences for each 1 Mb window across the genome. High 3D genome variability indicates a high average pairwise 3D divergence (i.e. diversity of 3D organization), while low 3D genome variability indicates low pairwise 3D divergence (i.e. similar 3D organization across all individuals).

#### Genomic windows with evidence of introgression

To define genomic regions with Neanderthal ancestry we used “segments” identified by Browning et al. [94] using Sprime, a heuristic scoring strategy that compares high-LD regions in a target admixed population (i.e., Europeans) with an unadmixed outgroup (i.e., Africans) to identify putatively introgressed regions. We considered a set of Sprime-identified segments shared (intersection) among East Asians (EAS), Europeans (EUR), and South Asians (SAS). We repeat the analysis using a more stringent subset of Sprime segments that (1) have at least 30 putatively introgressed variants that could be compared to the Altai Neanderthal genome and (2) had a match rate of at least 30% to the Altai Neanderthal allele (Neanderthal filter). We also considered the introgressed Neanderthal haplotypes previously identified by Vernot et al. [15] identified using the S* statistic. Finally, we consider introgressed segments unique to a single population (EAS, EUR, or SAS). Because these introgression calls only consider autosomes, we do not use the X chromosome for these analyses. Results from these sets of Neanderthal ancestry are in Figs. 5,S12,S13 and Tables S7,S8.

In the main text (Fig. 5), we compare the 3D genome variability between 1 Mb windows with no introgression (0%) versus windows where at least 70% of the bp have evidence of introgression. Other thresholds are shown in Fig. S12.

#### Predicting the amount of introgression

To test if 3D genome variability can be uniquely informative to predict tolerance of introgression, we conducted a simple linear regression. We predict the amount of introgression in a 1 Mb window while conditioning on the amount of sequence variability in a window. *Y* = *B*_0_ + *B*_1_*X*_3D variability_ + *B*_2_*X*_Sequence Variability_, where Y is the proportion of the 1 Mb window with evidence of introgression defined using the previously described sets of Neanderthal ancestry. For comparison, we also conducted some regressions where *Y* was modeled from only 3D variability or sequence variability alone. Results from these models are in Figs. 5B,S13, Tables S7,S8.

### 4.9 Individual-level introgression calls

We used introgression calls in 1KGP individuals from Chen et al. [96], which applied IBDmix with the Altai Neanderthal genome to identify introgressed segments in MHs. We identified windows with AH-MH divergence with evidence of introgression by intersecting with the introgression calls.

We also test the relationship between the amount of introgression an individual has and their 3D divergence from AHs. For each window, we compare the amount of introgression (% of bp) for an individual in a 1 Mb window with that individual’s 3D divergence from Neanderthals. We do this analysis for 15 Eurasians across 4,749 1 Mb autosomal windows (total *n* = 71, 235). In Fig. 6C we compare Eurasians to the Vindija Neanderthal 3D genome and in Fig. S16 we compare to Altai and Chagyrskaya. We also repeat the analysis removing windows with no evidence (0% bp) of introgression.

### 4.10 High-throughput chromosome conformation capture (Hi-C) analyses

#### 4.10.1 Experimental methods

Hi-C experiments of the three cell lines (HG00318A, HG01982 and GM12763C) were conducted following the manufacturer’s protocol of Arima-HiC Kit (A510008, Arima Genomics). Approximately 1 million cells from each cell line were counted and collected by centrifuging at 1100rpm for 5 min. Cells were subsequently fixed with 2% formaldehyde solution (252549, Sigma) rotating at room temperature for 20min and then quenched by adding 2.5M glycine to a final concentration of 0.2M rotating at room temperature for 10min. Cells were lysed and then digested using restriction enzyme. End-repair and biotin-label with biotin-14-dATP were performed followed by DNA ligation. DNA was then de-crosslinked and purified using 0.45X KAPA Pure Beads (KK8002, Roche). Quality control was performed following the Arima-QC1 Quality Control protocol. DNA was then sonicated into approximately 400bp fragments. Biotin-labeled DNA was enriched, end-repaired, adaptor ligated, and PCR amplified. KAPA Library Quantification Kit (KAPA Biosystems) was used to construct the Hi-C libraries based on manufacturer’s protocol.

Raw Hi-C data were first trimmed using Trim Galore (https://www.bioinformatics.babraham.ac.uk/projects/trim_galore/). Subsequently, the trimmed fastq files were processed using runHiC pipeline [111], which is a Hi-C data processing software and contains mapping, filtering and binning steps. First, reads were mapped to GRCh38 reference genome using bwa [112]. Note that for plotting comparisons to the archaic genomes, visualization is all in hg19 even though mapping was with GRCh38. Redundant PCR artifacts and read pairs that map to the same restriction fragment were removed. Multi-resolution cool (mcool) file, the official Hi-C data format for the 4DN consortium, was generated for each sample. Visualizations in this paper are using the 5kb resolution and visualized with adaptive coarsegrain smoothing [113]. We utilized Juicer tools [114] to generate multi-resolution.hic files for each sample.

#### 4.10.2 Quantifying differences in structure

We identified the subset of contacts which were *a priori* predicted to be most divergent using the computationally generated maps. To do this we took the top 0.1% (*n* = 100) of contacts in the computationally 3D matrix of the difference between Neanderthal counts versus African counts (*AH* −*MH*, Fig. S15B-C). This created a set of candidate loci for the contacts that were predicted to be the most strong in AH compared to MH. The experimental Hi-C maps are in 5 kB resolution while the computational maps are in 2048 bp bins so the candidate loci was interpolated to match the size of the experimental maps. The candidate loci were compared between the 3 Hi-C maps and plotted in Figs. 6C,S15D.

We also compared the 3D contact profiles along the location of the CTCF site that was modified by introgressed variation. Figs. 6D,S15E-F plot the normalized experimental contacts (y-axis) at the coordinates of the CTCF site to show the change in contact at this loci specifically (subset from the full 3D matrix). Given the noise in the experimental data, Figs. 6D,S15F show these data smoothed using a 3 bin rolling average. Fig S15E shows the raw data without smoothing.

### 4.11 Allele frequency, eQTL, and PheWAS analysis

Allele frequencies come from 1KGP Phase 3 [87] and are visualized in Fig. 6E using the Geography of Genetic Variants Browser [115]. eQTL analysis and plots were generated using the Genotype-Tissue Expression (GTEx) Project (V8 release) Portal (lifted over to hg19) [42]. PheWAS results are from the GWAS Atlas and consider 4756 traits [95]. The p-value line in Fig. 6G represents a highly conservative Bonferroni-corrected *P* -value (1.05 × 10^−5^) for testing 4756 traits, many of which are correlated traits and GWASs in which the SNP was not available to be tested.

### 4.12 Examples

The examples visualized in Figs. 3,6 are annotated using the UCSC genome browser [110]. They were each manually zoomed to highlight the regions of interest. We use ENCODE open chromatin candidate cis-regulatory elements (cCREs) [109] to highlight promoters (promoter-like signature, pink) and enhancers (proximal [orange] and distal [yellow] enhancer-like signature) combined from all cell types downloaded from the UCSC table browser (lifted over to hg19) [106]. We use Transcription Factor (TF) ChIP-seq Clusters (130 cell types) from ENCODE 3 [116, 117] downloaded from UCSC table browser [106]. We show the motif sequence logo with reference to the positive strand of hg19.

## 4.13 Data analysis and figure generation

The datasets we generated are available in the GitHub repository “neanderthal-3d-genome” available here https://github.com/emcarthur/neanderthal-3D-genome/ which will be formally cited and versioned upon publication.

All genomic coordinates and analysis refer to Homo sapiens (human) genome assembly GRCh37 (hg19), unless otherwise specified. All *P* values are two-tailed, unless otherwise specified. All measures of central tendencies are means, unless otherwise specified. Data and statistical analyses were conducted using Python 3.6.10 (Anaconda distribution), Jupyter Notebook, BedTools v2.26, and PLINK 1.9 [118, 119]. Figure generation was significantly aided by Matplotlib, Seaborn, Inkscape, and cooltools [113, 120–122].

### 4.14 Data availability

The publicly available data used for analysis are available in the following repositories. MH genome vcfs are from 1KGP (ftp.1000genomes.ebi.ac.uk/vol1/ftp/data_collections/1000_genomes _project/release/20190312_biallelic_SNV_and_INDEL/[87]. Archaic genotypes are from the following repositories: Altai Neanderthal [1] (ftp.eva.mpg.de/neandertal/Vindija/VCF/Altai/), Denisova (ftp.eva.mpg.de/neandertal/Vindija/VCF/Denisova/) [4], Vindija Neanderthal [2] (ftp.eva.mpg.de/neandertal/Vindija/VCF/Vindija33.19/), and Chagyrskaya Neanderthal [3] (ftp.eva.mpg.de/neandertal/Chagyrskaya/VCF/). Introgressed variants and segments are from Sprime Version 1 (https://data.mendeley.com/datasets/y7hyt83vxr)[94]. An alternative set of introgressed variants and segments are from S* (https://drive.google.com/drive/folders/0B9Pc7_zItMCVWUp6bWtXc2xJVkk?resourcekey=0-Cj8G4QYndXQLVIGPoWKUjQ)[15]]. Individual level 1KGP introgression calls are from the Akey Lab (https://drive.google.com/drive/folders/1mDQaDFS-j22Eim5_y7LAsTTNt5GWsoow)[96].

CTCF motifs are from genome-wide motif scans v1.0 (https://resources.altius.org/∼jvierstra/projects/motif-clustering/releases/v1.0/, all models in the CTCF archetype motif cluster, lifted-over to hg19)[108], CTCF-bound open chromatin candidate cis-regulatory elements (cCREs) in the HFF cell type (https://screen.encodeproject.org/ > Downloads > by cell type > HFF-Myc male newborn originated from foreskin fibroblast, lifted-over to hg19)[109], TAD boundaries in the HFF cell type are from processed MicroC data available at the 4D nucleome data portal (https://data.4dnucleome.org/experiment-set-replicates/4DNES9X112GZ/, lifted-over to hg19)[105]. RefSeq genes, TF ChIP-seq Clusters, enhancer and promoter cCREs are downloaded from the UCSC Table Browser (https://genome.ucsc.edu/cgi-bin/hgTables)[106]. Gene ontology annotations are downloaded from Enrichr (https://maayanlab.cloud/Enrichr/#libraries)[91–93]. System-level groupings of disease ontology terms were aided by Gene ORGANizer annotations(http://geneorganizer.huji.ac.il/downloads/)[107]. eQTL data is from the GTEx Portal (https://www.gtexportal.org/, lifted-over to hg19)[42]. PheWAS results are from the GWAS Atlas (https://atlas.ctglab.nl/)[95].

### 4.15 Code availability

Akita is in the “basenji” GitHub repository available here https://github.com/calico/basenji/tree/master/manuscripts/akita [82]. The “neanderthal-3d-genome” GitHub repository (above) contains a Jupyter notebook with custom code used for data analysis and all figure generation.

## 4.16 Acknowledgements

The authors would like to thank Colby Tubbs, Mary Lauren Benton, Douglas Ruderfer, Colin Brand, and other members of the Capra and Pollard labs for helpful discussions and manuscript comments. This work was conducted in part using the resources of the Advanced Computing Center for Research and Education (ACCRE) at Vanderbilt University, Nashville, TN.

## 4.17 Funding sources

This work was supported by the National Institutes of Health (NIH) General Medical Sciences award R35GM127087 to JAC, NIH National Human Genome Research Institute award F30HG011200 to EM, and T32GM007347. GF is supported by R35 GM143116-01. The funding bodies had no role in the design of the study and collection, analysis, or interpretation of data, or in writing the manuscript. The content is solely the responsibility of the authors and does not necessarily represent the official views of the NIH.

### 4.18 Authors’ contributions

EM, DCR, ENG, GF, MP, KK, FY, KSP, JAC conceived and designed the work presented here. EM, DCR, QW, YC, JW conducted all the analyses and experiments. EM, DCR, QW, ENG, GF, FY, KSP, JAC interpreted the results, drafted the work, and substantively revised the manuscript. EM, DCR, QW, YC, JW, ENG, GF, MP, KK, FY, KSP, JAC have approved the submitted version and have agreed to be accountable for their contributions.

### 4.20 List of Supplementary materials

Supplementary Text

Fig. S1 – S16

Table S1 – S8

