## Supplemental Text, Figures, Tables for "Reconstructing the 3D genome organization of Neanderthals reveals that chromatin folding shaped phenotypic and sequence divergence"

### 5 Supplementary Information

#### 5.1 Supplementary Text

When evaluating the relationship between 3D genome variability and introgression (Results section 2.7:"3D genome organization constrained introgression in MHs"), we considered a variety of subsets of genomic windows to fully explore these results. We show that the maintext results (Fig. 5) replicate when using earlier introgressed Neanderthal haplotype predictions from Vernot et al. [15] and other thresholds (Figs. S12,S13). We also find that 3D genome variability is more strongly predictive of introgression shared among all three super-populations than an introgressed sequence unique to a single super-population (Table S7). We hypothesize this is because the maintenance of a haplotype across diverse populations indicates stronger tolerance of the AH 3D organization pattern in diverse human genomic contexts. Additionally, 3D variability is relatively more informative about the amount of introgression when only considering windows of the genome with any introgressed sequence present (Table S8). Thus, we hypothesize that in 1 Mb windows with strong purifying selection against a large-effect introgressed variant (e.g., a deleterious protein-coding variant), 3D genome variability is less relevant. Ultimately, the pressures shaping the landscape of introgression across the genome were multi-factorial, but we demonstrate that 3D genome organization likely played a role.

#### 5.2 Supplementary Figures

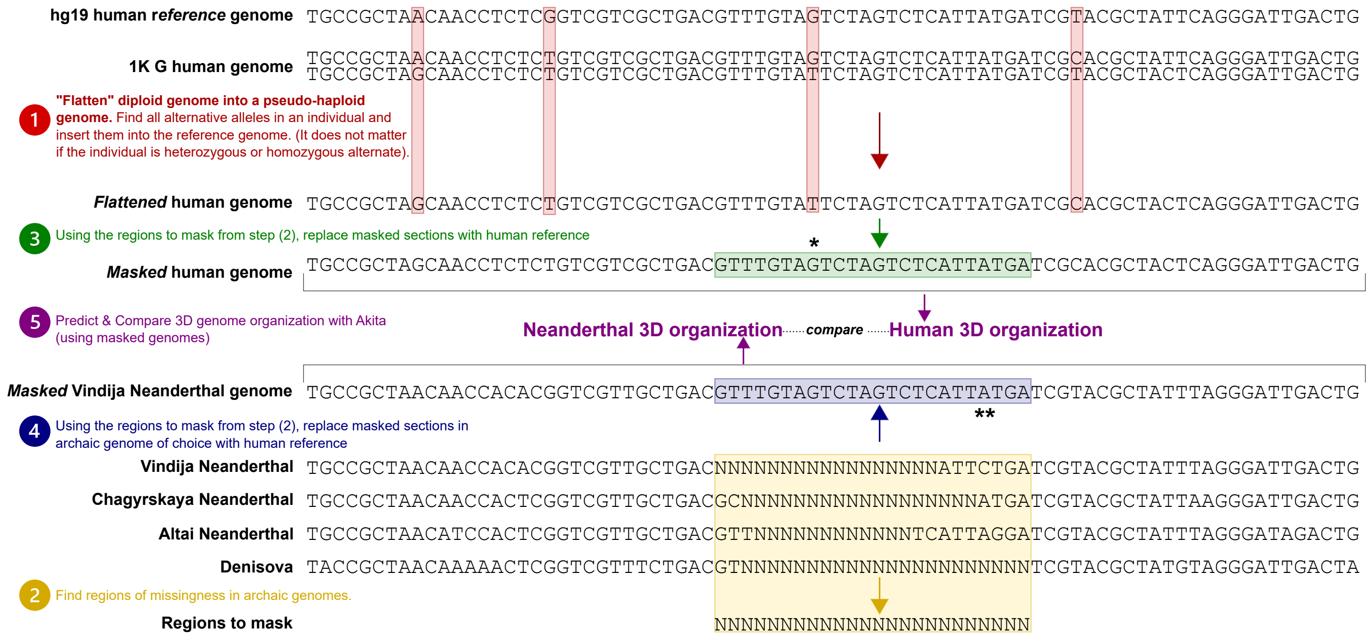


**Figure S1: Handling missingness in the archaic hominin genomes.** We constructed full-length genomes for each MH or AH based upon their genotyping information. Here, we illustrate a schematic of the procedure used to account for the challenges of archaic DNA. **(1)** Given the difficulty of distinguishing heterozygous genotypes in the ancient DNA samples, we treated all individuals as if they were homozygous (pseudo-haploid). If an individual had an alternate allele (homozygous or heterozygous), we inserted it into the reference genome to create a pseudo-haploid, or "flattened" genome for each individual (hightlighted in red boxes). **(2)** Because of gaps in coverage resulting from the challenges of ancient DNA, particularly in genomic regions of low complexity, we "masked" all genomic regions lacking archaic genotyping information by reverting nucleotide states to the hg19 reference (yellow box). For analyses that compared 3D genome organization between MHs and AHs, and MHs we do this masking procedure for both **[3]** MHs (green box) and **[4]** AHs (blue box) to facilitate appropriate comparisons. **[5]** We run Akita on each processed genome separately and then compare the resulting contact maps. By filling both genomes with the same sequence, there will be no differences between the AH-MH predictions or resulting comparisons. Although AHs and MHs certainly did not have the same genome sequences in these regions of missingness, we preferred this as a conservative approach to minimize identifying regions of interest if there were missing data. For example, we illustrate that at the nucleotide *****, although we observe an MH alternative allele (`T`), it gets masked and replaced with the hg19 reference (`G`) because that locus is not comparable to AH genomes. Many of the regions of missingness are shared by all or most of the AHs because those regions are just inherently difficult to sequence (Fig. S2). However, at the nucleotide ******, we illustrate another example where an allele observed in the Vindija genome (`C`) is masked with hg19 reference (`A`) so that it facilitates comparisons between the AHs (some of which have missingness at that locus).

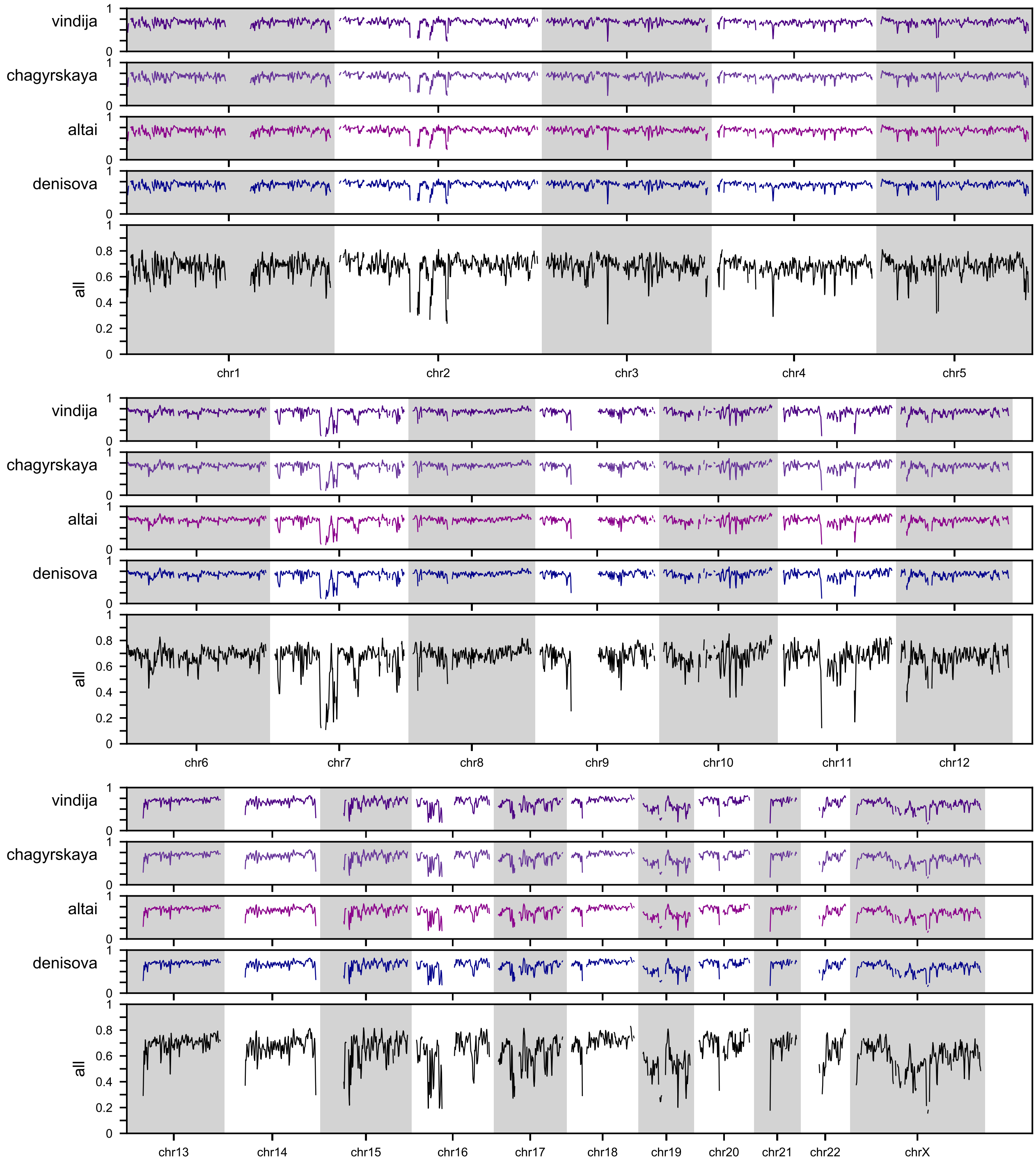


**Figure S2: Archaic hominin sequence coverage across the genome.** Ancient DNA fragmentation and degradation present challenges to both sequencing and alignment resulting in gaps in coverage, particularly in genomic regions of low complexity. Here, we show coverage across the genome for the 4 AHs. The horizontal axis represents genomic loci at the same sliding approximately 1 Mb window resolution ($N = 4,999$) used to do all analyses (Methods). The vertical axis unit is the proportion of bp with coverage (for the 1 Mb window). Bins without full coverage in modern humans (often near centromeres or telomeres) are excluded from all analyses and this figure. The bottom trace (black, labeled "all") represents the union of the missing segments for all 4 AHs. These regions are masked (Methods, Fig. S1) to facilitate 3D genome and sequence variation comparisons.

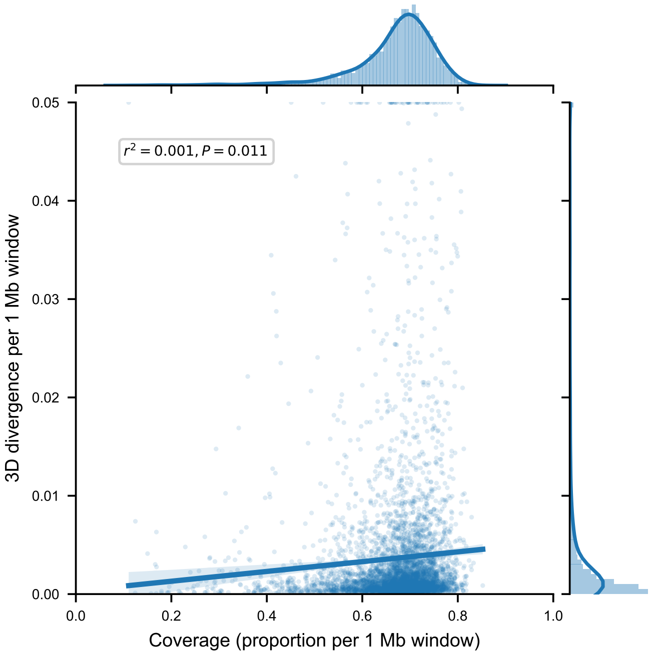


**Figure S3: 3D divergence in 1 Mb genomic window is weakly correlated with coverage.** Because we mask archaic missingness (Methods, Fig. S1,S2), regions with less coverage have more masking and the resulting processed sequences may have less AH-MH sequence variation. For 1 Mb windows across the genome ($N = 4999$), we compare AH (Vindija Neanderthal) and African MH (HG03105) 3D divergence (vertical axis) with the amount of coverage in that window (horizontal axis). The amount masked is equal to $1 - \text{coverage}$. 3D divergence is positively correlated with coverage ($r^2 = 0.001$, $P = 0.01$). This is likely because there is more opportunity to find variation that results in contact map changes when less of the region is masked; however, this correlation is very weak suggesting that more coverage of the archaic genomes may not uncover many additional examples of divergent organization.

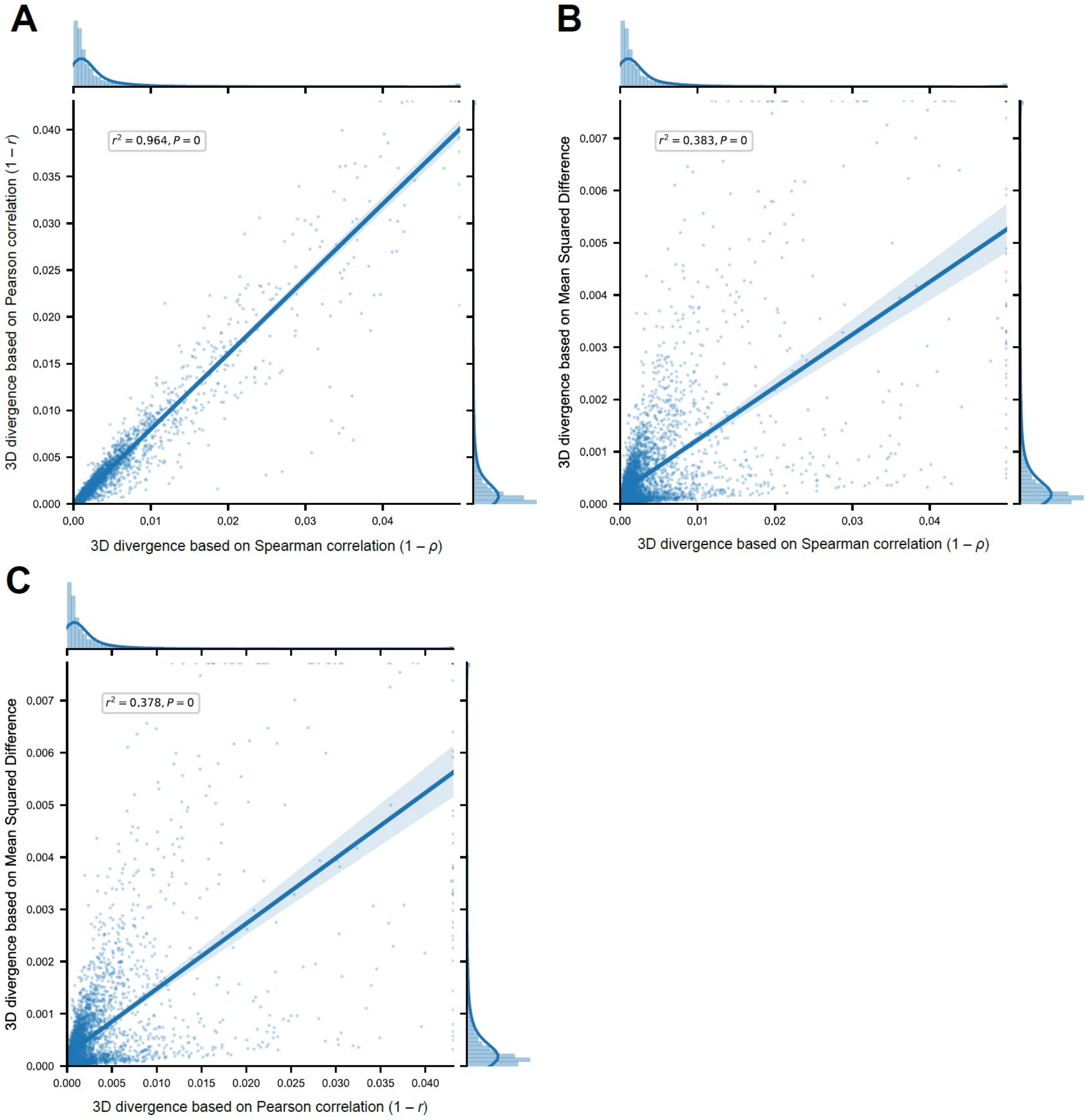


**Figure S4: Alternative measures of contact map comparison correlate with the 3D divergence derived from the Spearman's rank correlation coefficient.** In the main text, we compare chromatin contact maps using a 3D divergence score based on Spearman's rank correlation coefficient $(1 - \rho)$. Here, for the same windows across the genome $(N = 4999)$, we compare AH (Vindija Neanderthal) and African MH (HG03105) predictions using this Spearman-derived 3D divergence to others based on **(A)** Pearson's correlation coefficient $(1 - r)$ $(r^2 = 0.964)$ and **(B)** mean squared difference $(\frac{1}{n}\sum_{i=1}^{n}(x_i - y_i)^2)$ $(r^2 = 0.383)$. We also compare **(C)** these alternative measures (mean squared difference vs. Pearson's correlation) to each other $(r^2 = 0.378)$. The correlations between all measures are highly significant (all $P < 5 \times 10^{-324}$).

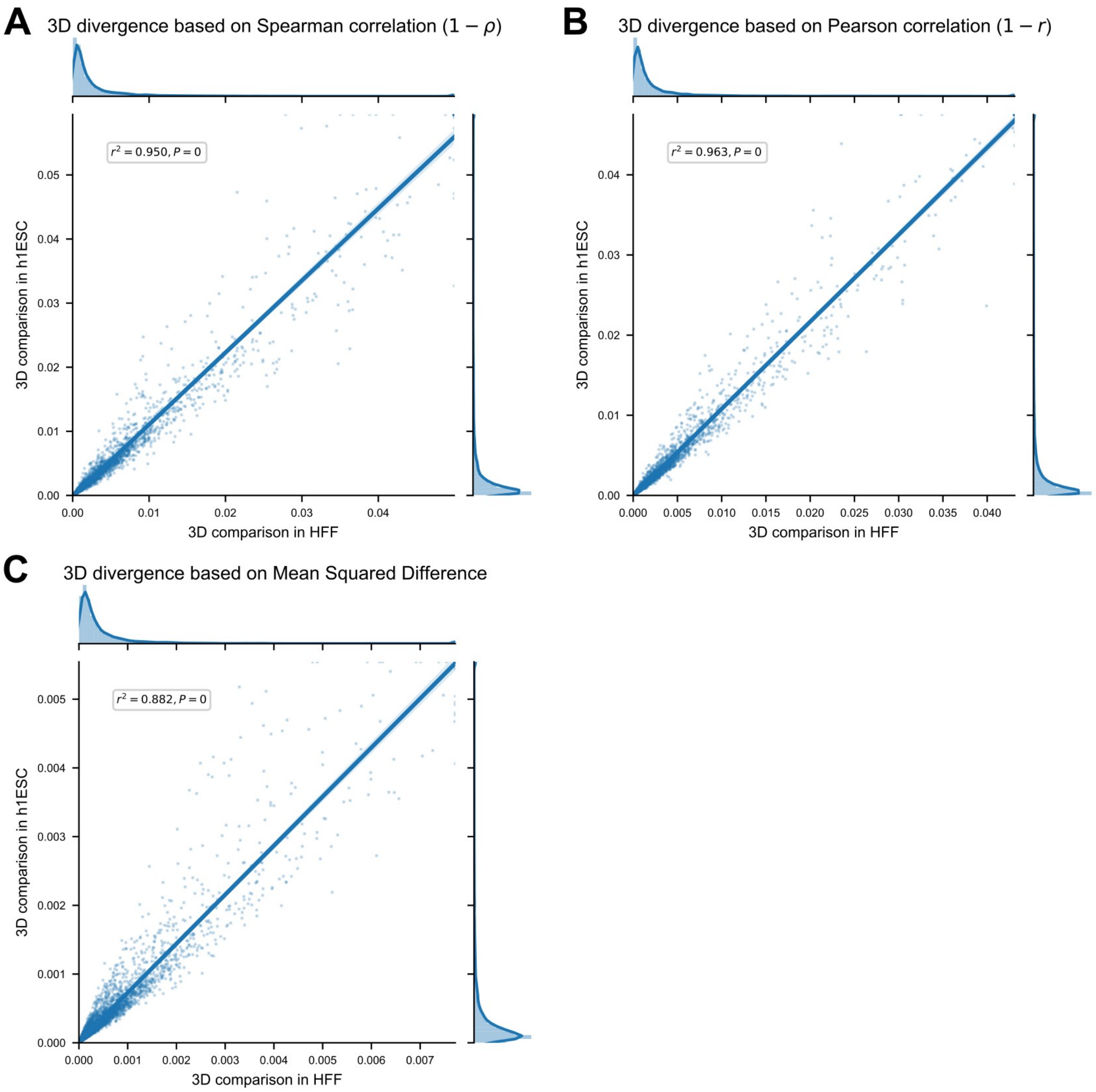


**Figure S5: 3D genome organization comparisons with chromatin contact maps from ESC are similar to those from HFF.** For the same windows across the genome ($N = 4999$), we compare AH (Vindija Neanderthal) and African MH (HG03105) predictions in embryonic stem cell (ESC) (vertical axis) versus human foreskin fibroblast (HFF) (horizontal axis) cell types. The comparisons across cell types are highly correlated regardless of the measure used to quantify their divergence. We consider comparison measures defined using the **(A)** Spearman correlation ($r^2 = 0.95$), **(B)** Pearson correlation ($r^2 = 0.96$), and **(C)** mean squared difference ($r^2 = 0.88$) (all $P < 5 \times 10^{-324}$).

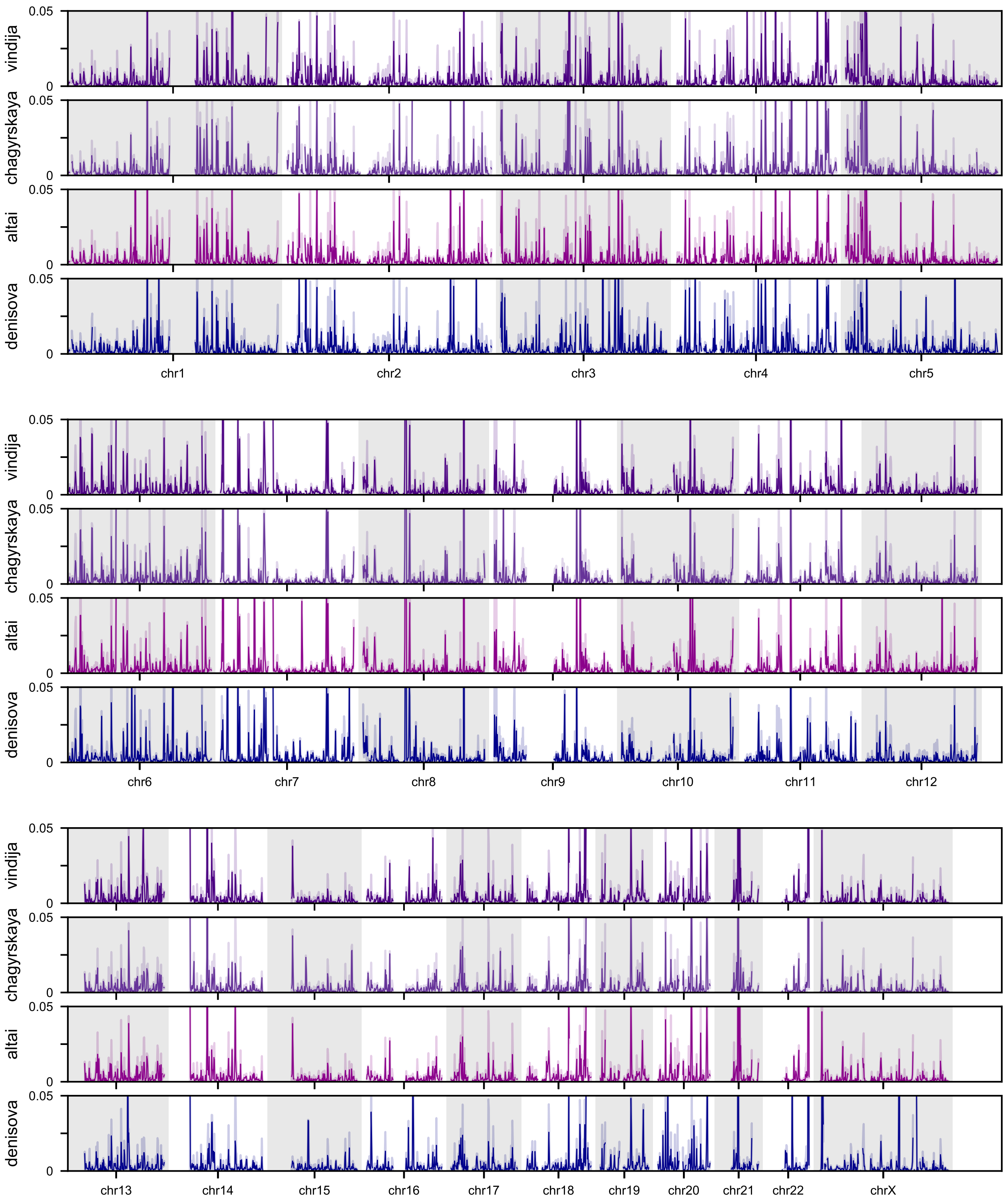


**Figure S6: AH-MH 3D divergence across the whole genome.** Across the genome, we plotted the average divergence of each of the AHs to five modern African individuals from different subpopulations. The horizontal axis represents genomic loci at the same sliding 1 Mb window resolution ($N = 4,999$) used to do all analyses (Methods). This expands Fig. 2C from chr7 to the whole genome. The error band indicates the 95% CI. Comparing the 3D genomes of Neanderthals (purple) or Denisova (blue) with MHs reveals windows of both similarity and divergence (peaks).

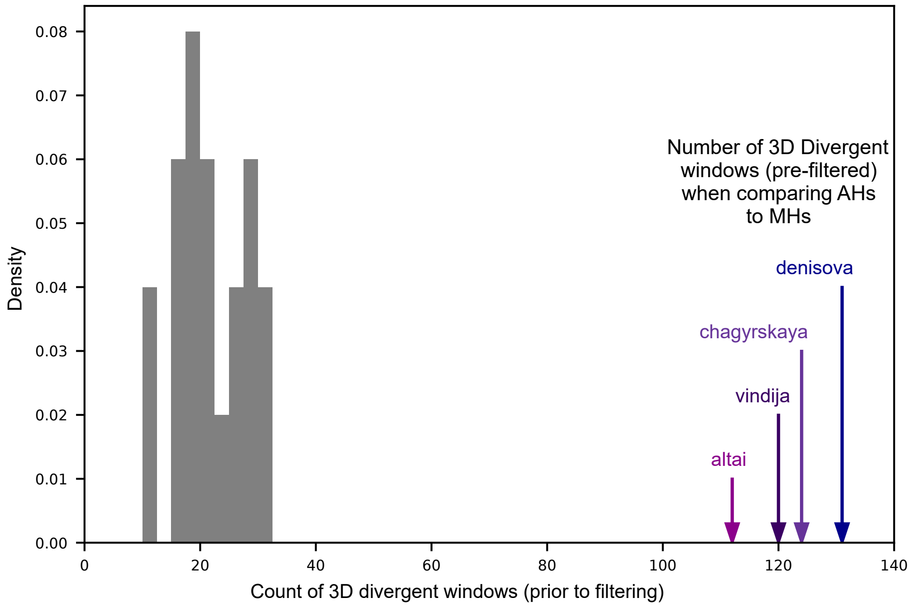


**Figure S7: When comparing archaic hominin genomes to African genomes, we observe many more 3D divergent windows than when comparing within the 20 African individuals.** To compare the number of divergent windows observed when comparing AHs to MHs (Fig. 2C), we use the same methods and thresholds to compare and call 3D divergent windows between 20 modern Africans. We compare each African 3D genome to the genomes of the other 19 African 3D genomes. Prior to filtering windows that are overlapping or have archaic-specific variants, we identify between 10-30 3D divergent windows for African-African comparisons. As expected, this is significantly less than the number of windows called when comparing AHs to the African individuals to call 3D divergent regions of the genome.

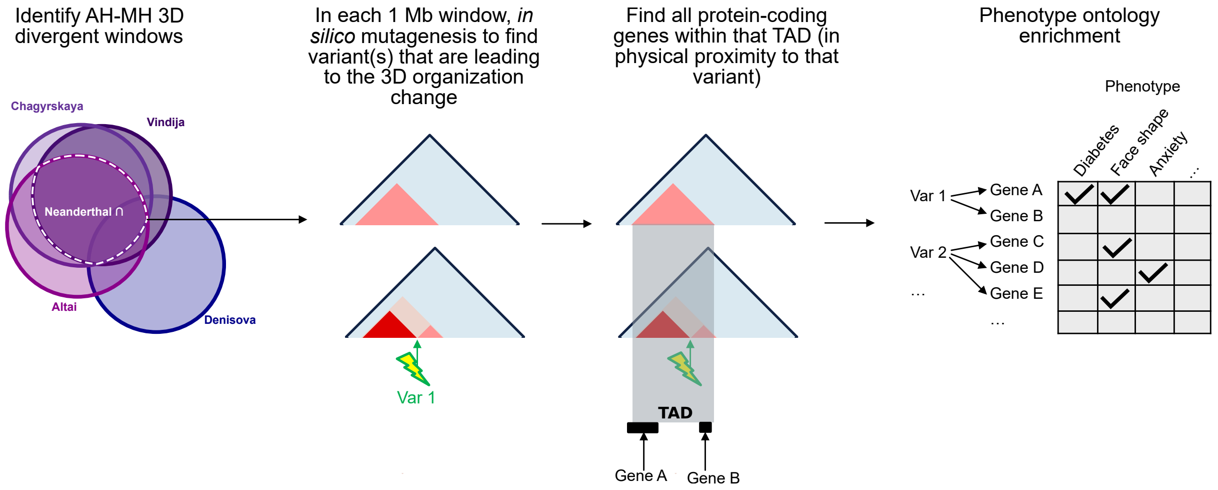


**Figure S8: Method for linking 3D divergent windows to test phenotype ontology term enrichment.** To test if differences in AH-MH 3D organization are enriched near genes related to particular phenotypes we follow a procedure that sequentially links 3D divergent windows to variants to TADs to genes and, ultimately, to phenotypes. We identify AH-MH 3D divergent windows in Fig. 3A–B. We consider three different sets of AH-MH divergent windows, those shared (intersect) by all Neanderthals, those in any Neanderthal (union), and those in Denisova. Results from the set shared by all Neanderthals ($N = 43$ windows) are shown in the main text (Fig. 3D). In each 1 Mb 3D divergent window, we identify the variant(s) contributing to the most prominent 3D differences using *in silico* mutagenesis (lightning bolt) (Methods). 3D-modifying variants are then linked to protein-coding genes (black bars) in their TAD (gray rectangle) because this provides evidence of physical proximity. Genes are linked to phenotypes from the Human Phenotype Ontology (HPO) and genome-wide association studies (GWAS) Catalog 2019. Through this procedure, we counted the number of ontology terms linked to the set of 3D-modifying variants. We test enrichment for ontology terms linked to at least one 3D-modifying variant using a shuffling approach to create an empirical distribution for how many times we would observe each annotation under the null. We used these distributions to calculate an enrichment and $P$-value for each ontology term. The specific data sets used in this procedure are detailed in the Methods. Counts of the number of windows, 3D-modifying variants, genes, and phenotypes for each set are in Table S2. Results for enrichment are in Figs. 3D,S9.

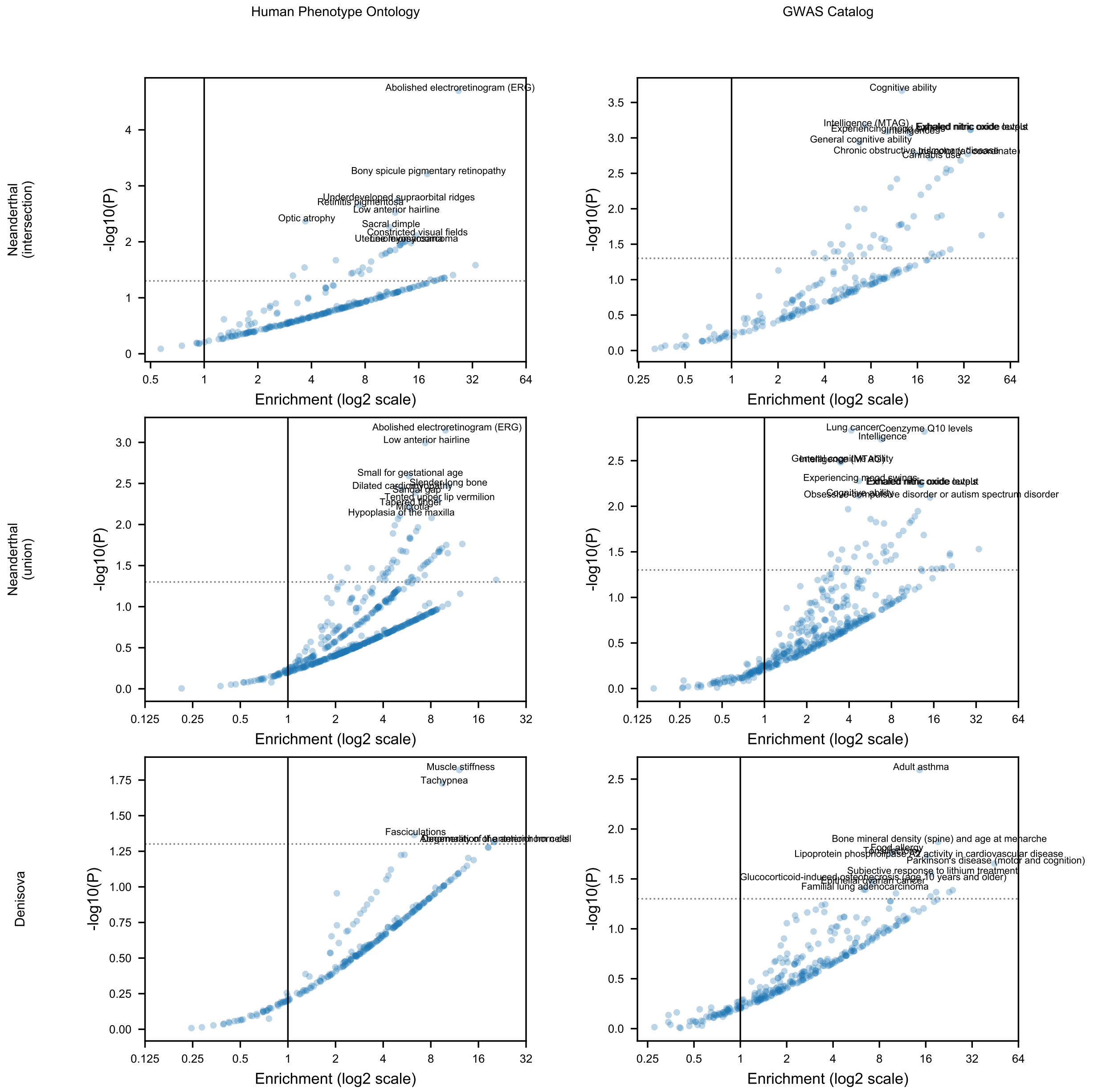


**Figure S9: Phenotype ontology enrichment across other sets of AH-MH 3D divergent windows implicate similar phenotypes.** When testing if differences in AH-MH 3D organization are enriched near genes related to particular phenotypes, we used three different sets of AH-MH 3D divergent windows (rows) and two different sets of gene-phenotype links (columns). The top set is from 43 3D-divergent windows shared by Neanderthals (intersect) (also shown in the main text, Fig. 3D). The middle is from 110 divergent windows in any Neanderthal (union). The bottom is from 73 divergent windows in Denisova. Each volcano plot has enrichment on the horizontal axis and significance on the vertical axis which were calculated with reference to a shuffled null distribution ($n = 500,000$, Methods). Each point represents one ontology term. Only terms linked to the 3D divergent windows in each set were tested for enrichment or depletion. Terms that were not linked to any 3D divergent window are depleted and are therefore not plotted or tested (hence why most terms with any overlap are positively enriched rather than depleted). The most significant 10 terms are labeled if $P < 0.05$ (dotted line). Similar to the Neanderthal (intersection) set, phenotypes related to the retina, hair, immune response, skeleton, cognition, and lung capacity are highlighted. Additional phenotypes at nominal significance include traits related to the heart, muscle, cancer, and bone density. Details about the process to link the 3D divergent windows to genes and phenotypes are in the Methods and Fig. S8. Details about the number of windows, variants, and phenotypes considered for each set are in Table S2.

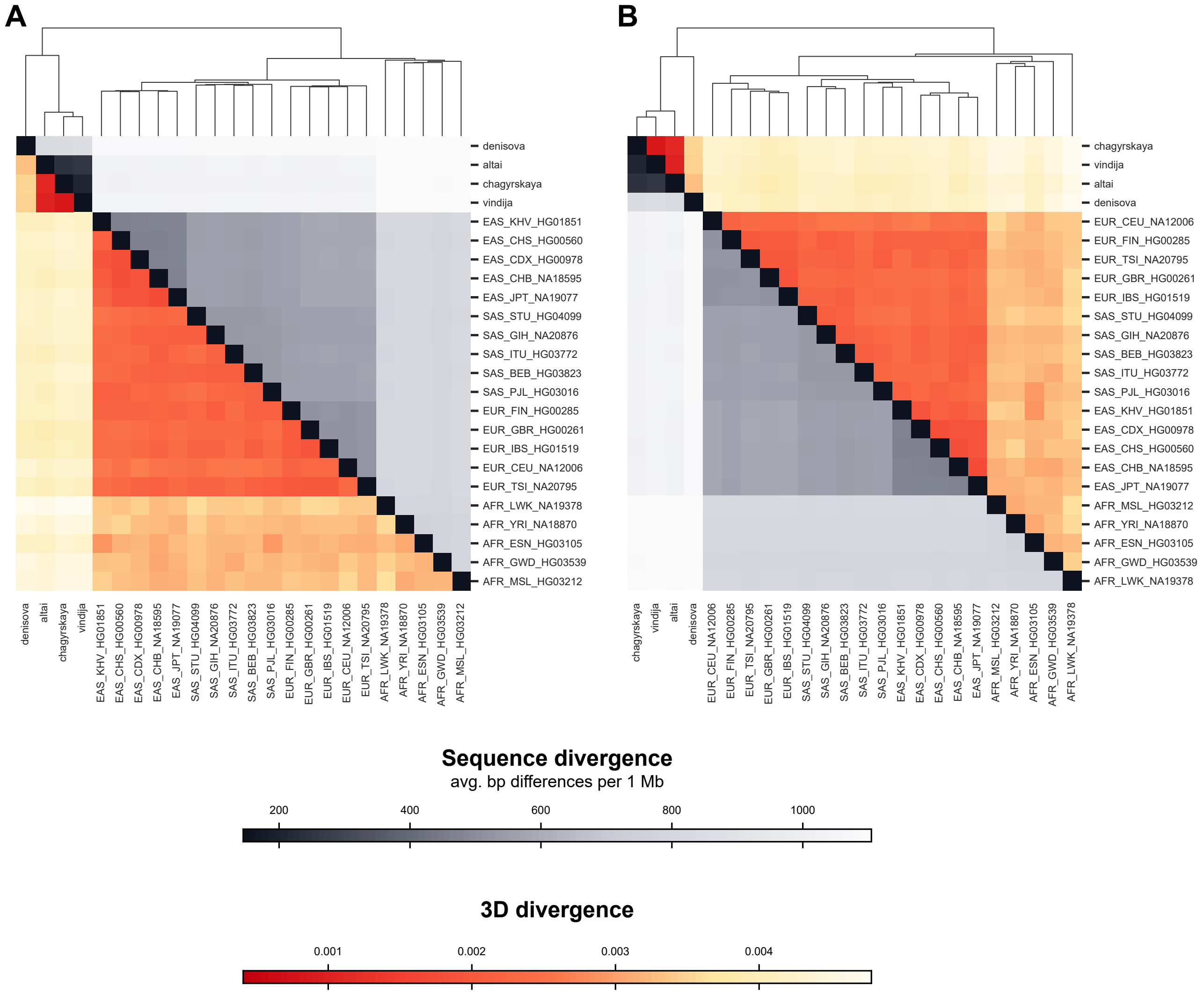


**Figure S10: Full pairwise heatmaps clustered by both sequence 3D divergence and sequence divergence.** We calculated the mean genome-wide 3D divergence for all pairs of AH and MH individuals (oranges) to compare with the genome-wide mean sequence divergence (grays). Fig. 4A displays these heatmaps when clustered by sequence divergence. Fig. 4A is reproduced in **(A)** with the full labels of all 1KGP individuals and their sub- and super-population information. **(B)** We also show the heatmap clustered by 3D genome divergence. Overall, global patterns of 3D genome divergence follow global patterns of sequence divergence. Lists of 1KGP individuals used and their abbreviation codes are defined in Table S1.

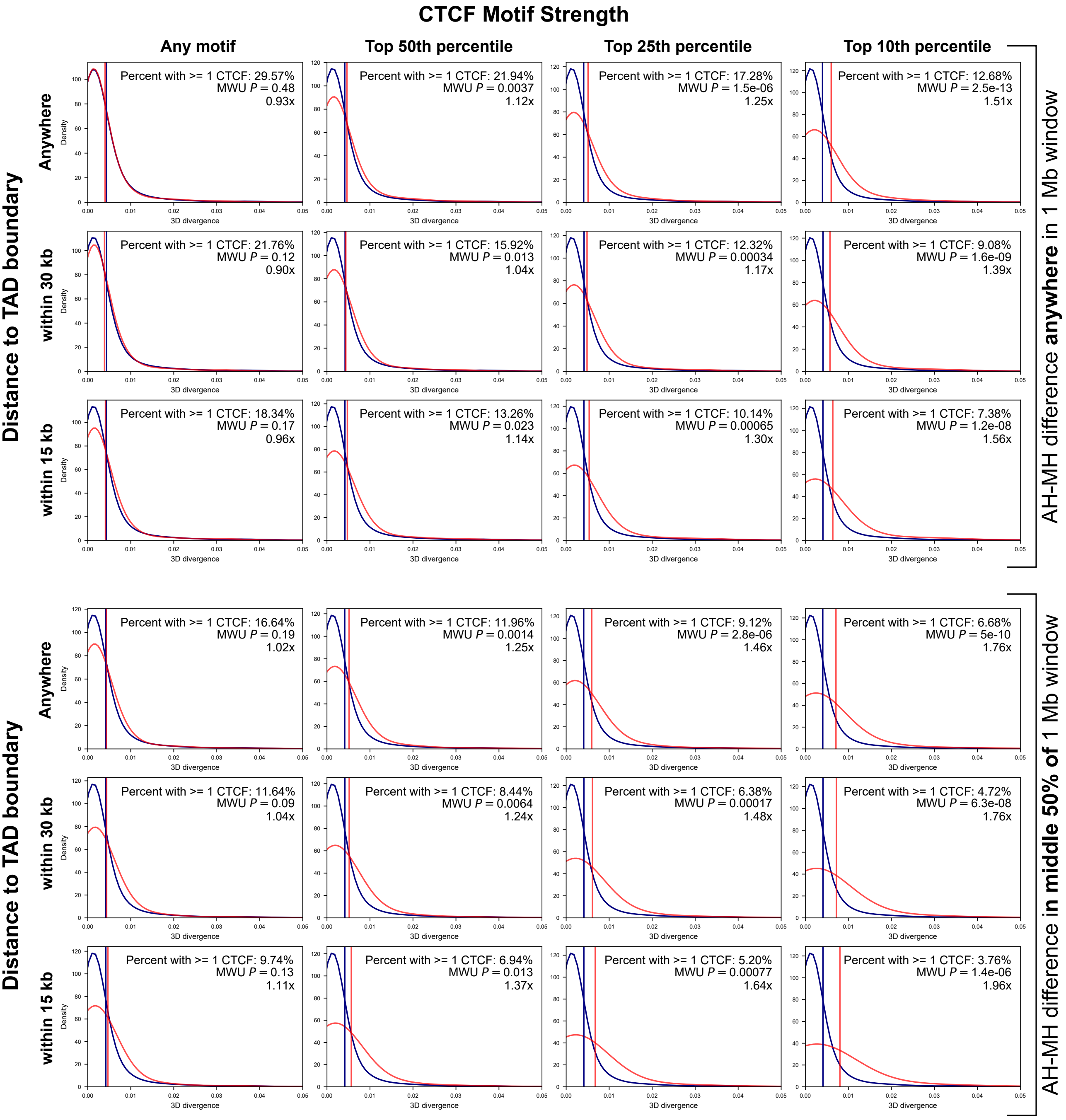


**Figure S11: 3D genome divergence depends on both the strength and context of the CTCF motif disrupted.** Based on the importance of CTCF-binding in maintaining 3D genome organization, we quantified the effects of AH-MH nucleotide differences overlapping CTCF binding motifs on 3D divergence. Given the complexity in the "grammar" of encoding 3D genome organization, we hypothesized that not all CTCF disruptions are equally likely to influence 3D divergence. Fig. 4B demonstrates this. But, here we replicate this with other thresholds and filters. We considered if each 1 Mb window ($N = 4,999$) had a sequence difference between a Neanderthal (Vindija) and a MH (HG03105) genome that overlapped a CTCF site. We plotted the distribution of 3D divergence in a window by whether there was a "CTCF overlapping variant" (red) or not (blue). We further filtered windows by multiple annotations describing the context and strength of the CTCF site overlapped. First, we stratified windows by if the "CTCF overlapping variant" occurs within the middle half of the 1 Mb window (right vertical axis). Second, we stratified windows by the proximity of the "CTCF overlapping variant" to a TAD boundary (anywhere, within 30 kb, or within 15 kb) (left vertical axis). Finally, we stratified windows by the strength of the overlapped CTCF motif in percentiles (any, top 50%, 25%, or 10%) (horizontal axis). All three features describing context and strength are informative about the likelihood of 3D divergence. For example, when filtering for the strongest CTCF motifs overlapped by a variant, 3D divergence increases 1.96-fold compared to 1.11-fold if strength is ignored (bottom left vs. bottom right). When considering by proximity to TAD boundaries, 3D divergence always increases when a "CTCF overlapping variant" is closer to a TAD boundary ($4^{\text{th}}$ row vs. $6^{\text{th}}$ row). This illustrates that our approach has learned the complex sequence patterns underlying 3D genome folding that could not be determined by simply intersecting AH variants with all CTCF sites.

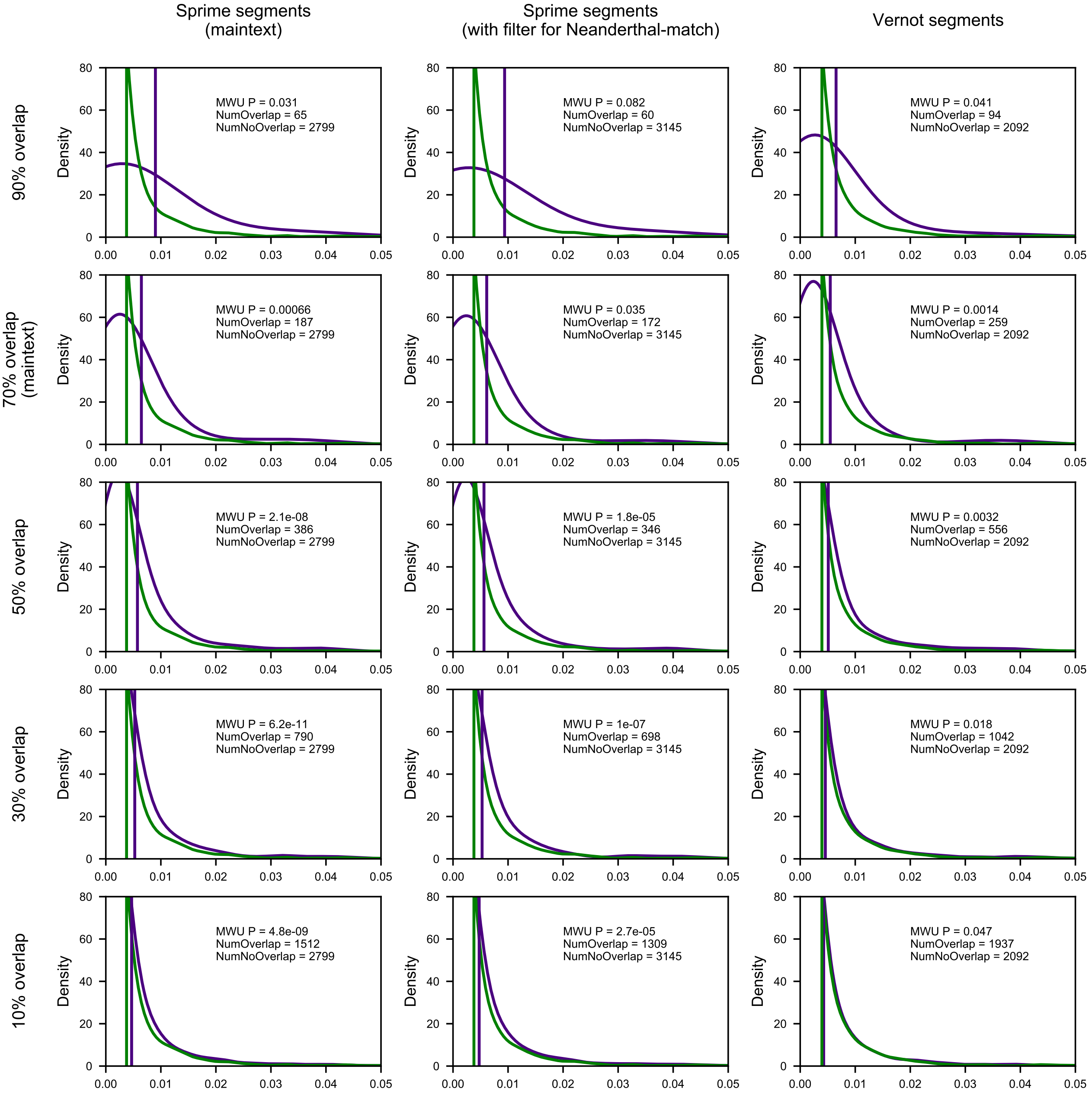


**Figure S12: Windows with evidence of AH introgression are more 3D variable in MHs even when using different definitions of introgression.** Genomic windows with high levels of introgression across present-day non-African populations (purple distribution) are more 3D-variable in modern Africans (horizontal axis) than windows without evidence of introgression (green distribution). In the main text, we considered introgression defined by segments from Browning et al. [94] (first column) covering at least 70% of bases in a 1 Mb window (second row). This identifies 187 autosomal 1 Mb windows with introgression and 2,799 without (same figure as Fig. 5A). Here, we show that this trend is consistent even when using different sets of introgressed haplotypes (columns) and thresholds for overlap (rows). Sprime segments are from Browning et al. [94]. Sprime segments with Neanderthal-matching filter are a subset of the Browning et al. [94] introgressed segments that have 30 putatively introgressed variants that could be compared to the Altai Neanderthal genome and had a match rate of at least 30% to the Altai Neanderthal allele. S* Vernot segments are from Vernot et al. [15]. Vertical lines represent the distribution means. $P$-values are from a two-tailed Mann–Whitney U test.

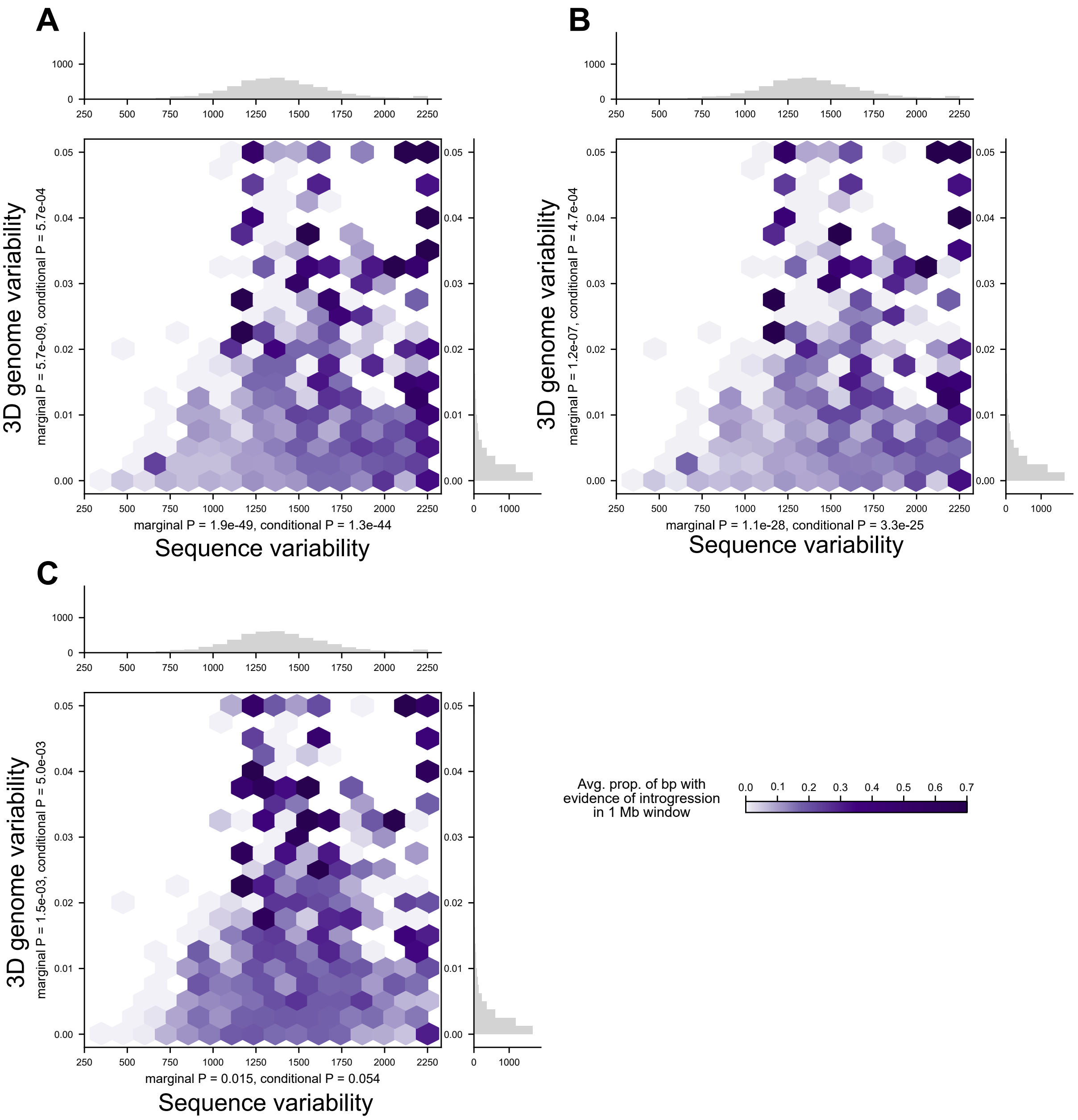


**Figure S13: 3D variable windows in MH have more evidence of AH introgression even when using different definitions of introgression.** For three different sets of introgressed haplotypes (**A**-**C**), we plot the relationship between sequence variability (horizontal axis) and 3D genome variability (vertical axis) with amount of AH ancestry in a window (purples). Darker purple indicates a higher proportion of introgression in a 1 Mb genomic window. 3D genome variability is defined as the average modern-African pairwise 3D genome diversity. Sequence variability is defined as the average pairwise nucleotide differences per modern-African in a 1 Mb window. *P*-values correspond to the significance of sequence variability or 3D genome variability to predict amount of introgression in a 1 Mb window. 3D genome variability is predictive of the amount of introgression both independently and when conditioned on sequence variability for all three sets of introgression. For, **A**,**B**, and **C**, respectively, introgressed haplotypes are from Sprime segments, Sprime segments with a Neanderthal-sequence match filter, and S* segments. **A** is shown in the maintext in Fig. 5B. Sprime segments are from Browning et al. [94]. Sprime segments with Neanderthal-matching filter are a subset of the Browning et al. [94] introgressed segments that have 30 putatively introgressed variants that could be compared to the Altai Neanderthal genome and had a match rate of at least 30% to the Altai Neanderthal allele. Vernot segments are from Vernot et al. [15].

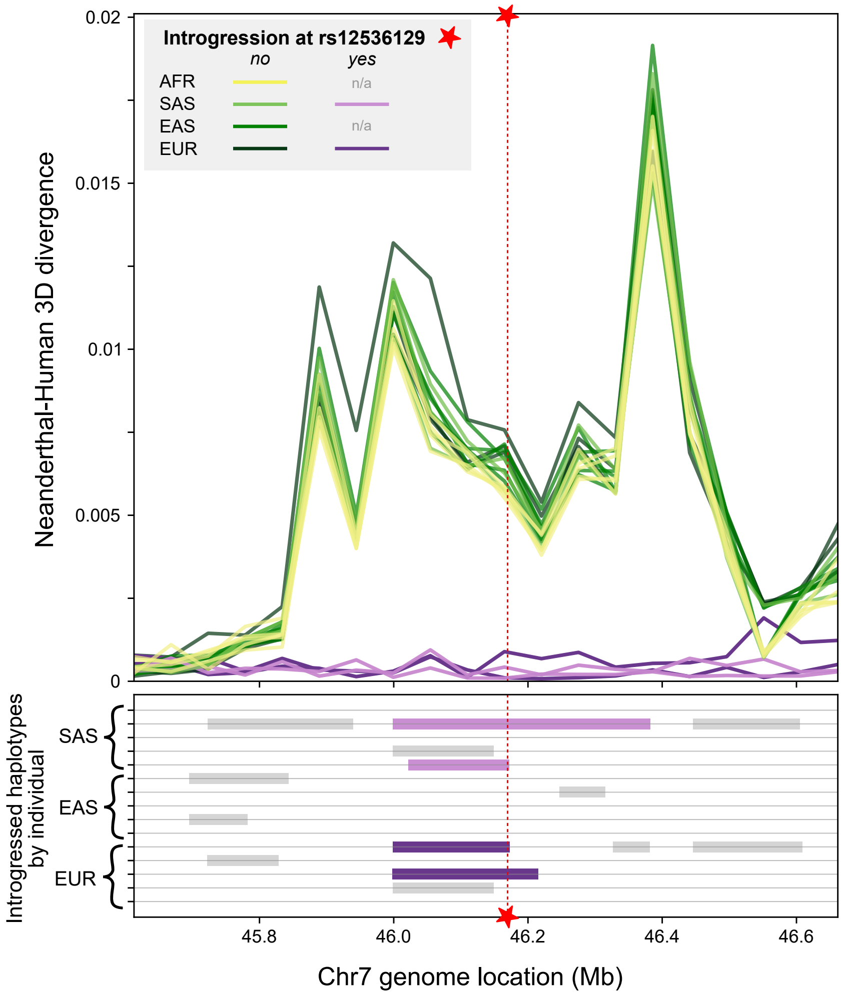


**Figure S14: Eurasians with introgression at chromosome 7 locus have 3D genome organization similar to Neanderthals.** Comparison of the 3D contact maps between Neanderthal (Vindija) and 20 MHs for a window on chromosome 7 reveals that most MHs (yellow, green) have different 3D organization compared to Neanderthals. In contrast, four MHs with introgression (purple boxes) overlapping chr7:46,169,621 (red star) have similar 3D organization compared to Neanderthals across this part of the genome (purple). Other individuals have introgression within the greater genomic window (gray boxes); however, the status of introgression at rs12536129 (red star) correlates with 3D genome divergence. This identifies a region where Eurasians with introgression inherited a novel 3D genome folding pattern from Neanderthal introgression. This locus is explored in Fig 6. (AFR: African, SAS: Southeast Asian, EAS: East Asian, EUR: European)

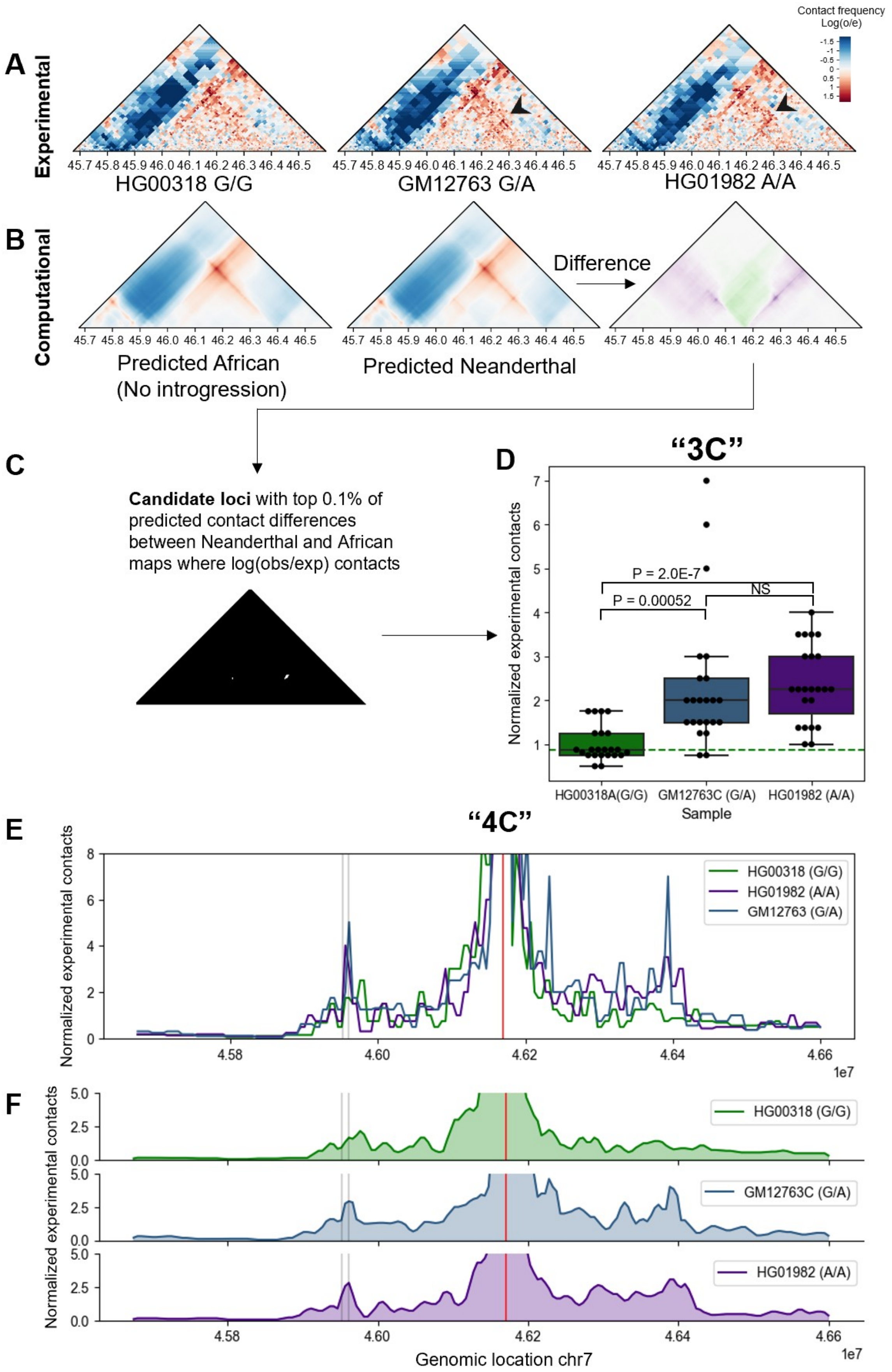


**Figure S15: Experimental Hi-C validates computational predicted introgressed 3D substructure at chromosome 7 locus.** Caption on next page.

**Figure S15: Experimental Hi-C validates computational predicted introgressed 3D substructure at chromosome 7 locus (A)** Experimental Hi-C maps for three individuals with differential introgression at the chr7 locus explored in Fig. 6 (chr7:45674599-46599598 shown here). The individual heterozygous (GM12763) for the introgressed allele at chr7:46169621 (rs12536129) demonstrates substructure similar to the individual homozygous for introgression (HG01982). **(B)** The difference between the computationally predicted 3D organization for African (ancestral) and Neanderthal (derived) is used to identify the top 0.1% (top 100 contacts or cells in the matrix) *a priori* depicted in **C**. **(D)** The individuals with introgression (GM12763, HG01982) have significantly increased 3D contacts at this locus. The individual homozygous for introgression (HG01982) has increased contacts when compared to the individual heterozygous for introgression (GM12763); however, not significantly so. We theorize that there may be a dominant effect of the increased strength of the CTCF motif. **(E,F)** 3D contact profiles along the location of the CTCF site that was modified by introgressed variation. **E** depicts the raw data and **F** show these data smoothed using a three-bin rolling average. The gray lines denote the location of *IGFBP3* and the red line is the location of the CTCF motif strengthened by introgression.

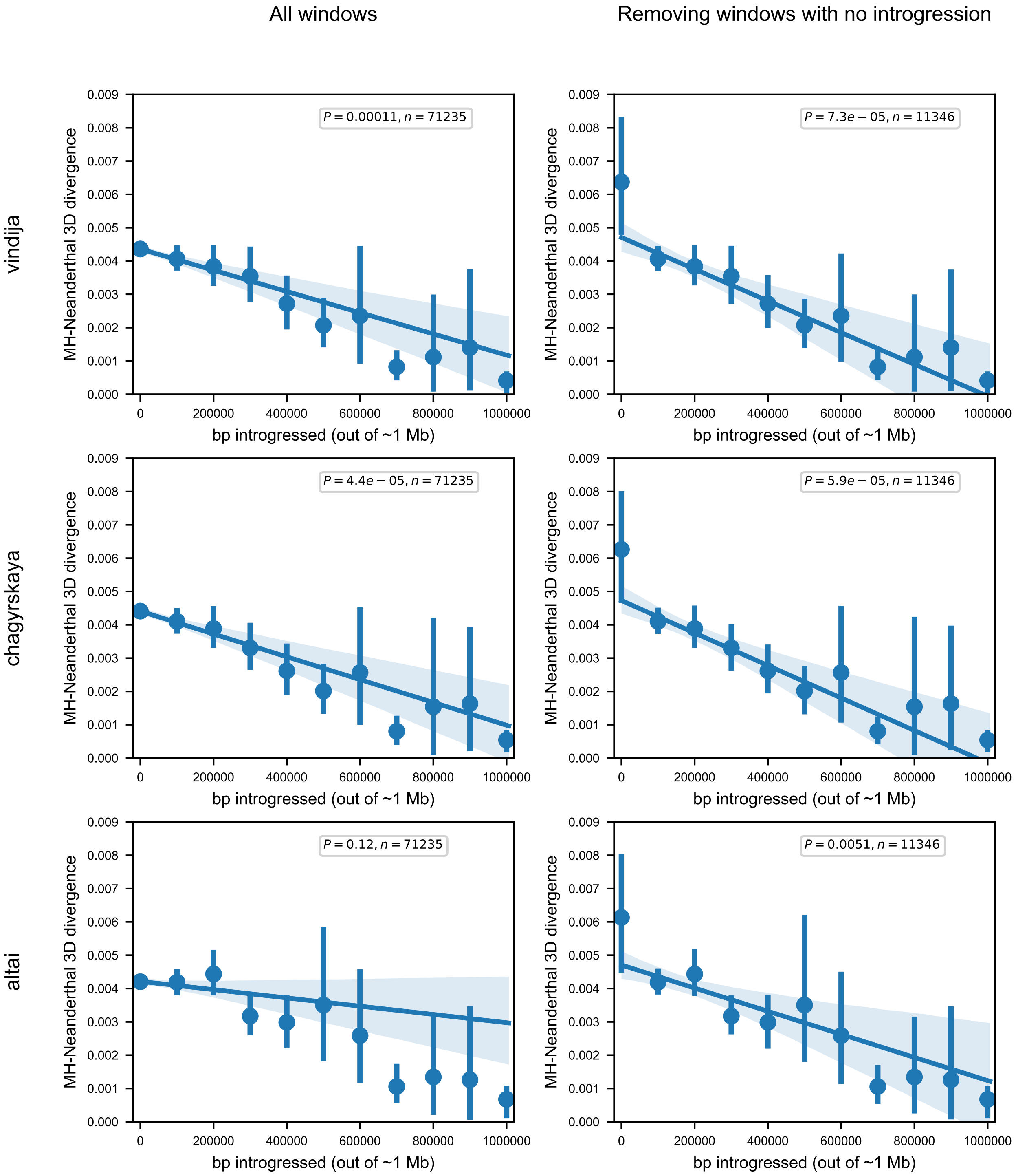


**Figure S16: Amount of introgression is negatively correlated with 3D divergence to all Neanderthal individuals.** The amount of introgression in a 1 Mb window (number of bp, horizontal axis) is significantly correlated with the similarity of an individual's 3D genome organization to a Neanderthal's genome organization (vertical axis). This is demonstrated across all three Neanderthal individuals: Vindija in the top panel (also shown in Fig. 6C), Chagyrskaya in the middle, and Altai at the bottom. We hypothesize the trend is weakest in Altai because it is less related to the introgressing Neanderthal population compared to the Vindija Neanderthal [2]. The left column considers all 4,749 autosomal 1 Mb windows for 15 Eurasians (total $n = 71,235$, 1KGP individuals in Table S1). In the right column, this trend also holds when you remove 1 Mb windows with no (0 bp) introgression in the 15 considered Eurasian individuals $n = 11,346$. The $P$-values are the significance of the correlation. The error bars signify 95% bootstrapped confidence intervals and the error band signifies the 95% bootstrapped confidence interval for the linear regression estimate.

#### 5.3 Supplementary Tables

| | Superpopulation | Subpopulation | ID | Subpopulation Description |
|---|---|---|---|---|
| Individuals in initial and Eurasian introgression analyses | EAS | CDX | HG00978 | Chinese Dai in Xishuangbanna, China |
| | EAS | CHB | NA18595 | Han Chinese in Beijing, China |
| | EAS | CHS | HG00560 | Han Chinese South |
| | EAS | JPT | NA19077 | Japanese in Tokyo, Japan |
| | EAS | KHV | HG01851 | Kinh in Ho Chi Minh City, Vietnam |
| | EUR | CEU | NA12006 | Utah residents (CEPH) with Northern and Western European ancestry |
| | EUR | FIN | HG00285 | Finnish in Finland |
| | EUR | GBR | HG00261 | British in England and Scotland |
| | EUR | IBS | HG01519 | Iberian populations in Spain |
| | EUR | TSI | NA20795 | Toscani in Italia |
| | SAS | BEB | HG03823 | Bengali in Bangladesh |
| | SAS | GIH | NA20876 | Gujarati Indian in Houston, TX |
| | SAS | ITU | HG03772 | Indian Telugu in the UK |
| | SAS | PJL | HG03016 | Punjabi in Lahore, Pakistan |
| | SAS | STU | HG04099 | Sri Lankan Tamil in the UK |
| | AFR | GWD | HG03539 | Gambian in Western Division, The Gambia |
| | AFR | LWK | NA19378 | Luhya in Webuye, Kenya |
| | AFR | MSL | HG03212 | Mende in Sierra Leone |
| | AFR | YRI | NA18870 | Yoruba in Ibadan, Nigeria |
| | AFR | ESN | HG03105* | Esan in Nigeria |
| Africans in AH-MH divergence and 3D genome variability analyses | AFR | ESN | HG03105 | Esan in Nigeria |
| | AFR | ESN | HG03499 | Esan in Nigeria |
| | AFR | ESN | HG03511 | Esan in Nigeria |
| | AFR | ESN | HG03514 | Esan in Nigeria |
| | AFR | ESN | HG02922 | Esan in Nigeria |
| | AFR | GWD | HG03539 | Gambian in Western Division, The Gambia |
| | AFR | GWD | HG03025 | Gambian in Western Division, The Gambia |
| | AFR | GWD | HG03028 | Gambian in Western Division, The Gambia |
| | AFR | GWD | HG03040 | Gambian in Western Division, The Gambia |
| | AFR | GWD | HG03046 | Gambian in Western Division, The Gambia |
| | AFR | LWK | NA19378 | Luhya in Webuye, Kenya |
| | AFR | LWK | NA19017 | Luhya in Webuye, Kenya |
| | AFR | LWK | NA19434 | Luhya in Webuye, Kenya |
| | AFR | LWK | NA19445 | Luhya in Webuye, Kenya |
| | AFR | LWK | NA19019 | Luhya in Webuye, Kenya |
| | AFR | MSL | HG03212 | Mende in Sierra Leone |
| | AFR | MSL | HG03086 | Mende in Sierra Leone |
| | AFR | MSL | HG03085 | Mende in Sierra Leone |
| | AFR | MSL | HG03437 | Mende in Sierra Leone |
| | AFR | MSL | HG03378 | Mende in Sierra Leone |

**Table S1: 1KGP individual genomes used for 3D genome predictions.** The top set of individuals were used in the initial 3D genome survey (Figs. 2, 4A) and introgression analyses (Fig. 6). The bottom set of African individuals was used to more robustly call AH-MH 3D genome divergence windows (Fig. 3) and to calculate MH 3D genome variability (Fig. 5). For consistency, the genome of HG03105 was used for all examples.

| ‘ | Number of 1 Mb 3D divergent windows (includes partially overlapped windows) | Number of unique 3D divergent windows (merging overlapping windows) | Number of “3D-modifying variants” observed in the 3D divergent windows | Number of “3D-modifying variants” that have evidence of introgression | Proportion of “3D-modifying variants” that have evidence of introgression | Number of gene-”3D-modifying variants” links* | Number of unique genes linked to 3D-modifying variants”* | Number of unique HPO terms linked to “3D-modifying variants” (used in enrichment test)* | Number of unique GWAS terms linked to “3D-modifying variants” (used in enrichment test)* |
|---|---|---|---|---|---|---|---|---|---|
| **Vindija** | 93 | 70 | 76 | 38 | 0.500 | | | | |
| **Chagyrskaya** | 95 | 71 | 78 | 32 | 0.410 | | | | |
| **Altai** | 82 | 67 | 73 | 33 | 0.452 | | | | |
| **Denisova** | 105 | 73 | 83 | 9 | 0.108 | 130 | 129 | 248 | 318 |
| **All Neanderthals (intersection)** | 54 | 43 | 45 | 28 | 0.622 | 88 | 85 | 271 | 208 |
| **All Neanderthals (union)** | 144 | 110 | 121 | 43 | 0.355 | 224 | 206 | 535 | 435 |
| **All archaic hominins (intersection)** | 10 | 7 | 8 | 6 | 0.750 | | | | |
| **All archaic hominins (union)** | 234 | 167 | 191 | 45 | 0.236 | | | | |

**Table S2: Counts of 3D divergent windows and 3D-modifying variants.** The number of 3D divergent windows per different AH individuals (rows) are in the first two columns. The first column is the raw 1 Mb windows, while the second column counts overlapping windows as one merged window (these values are depicted in Fig. 3B). The number of 3D-modifying variants in each window is in column three. The number and fraction of these 3D-modifying variants that are introgressed are in columns four and five. To conduct the phenotype ontology enrichment analyses shown in Figs. 3D,S9, we linked the 3D-modifying variants to genes (column six and seven, see Methods). These genes were then linked to terms using HPO (column eight) and the GWAS Catalog. These two sets of terms were then tested for enrichment. The phenotype ontology enrichment analysis parts (columns denoted with *) were only calculated on certain sets of 3D-diverged windows.

See supplementary excel file for large tables.

**Table S3: AH-MH 3D divergent windows.** Coordinates (in hg19) for 167 AH-MH 3D divergent windows identified in Fig. 3A–B. Windows were identified at approximately 1 Mb resolution (Methods) and overlapping windows were merged. The "AH" column details for which AH(s) the 3D divergent window was identified (A: Altai, C: Chagyrskaya, D: Denisova, V: Vindija). If a window was identified in two different AHs but they were only partially overlapping, the specific coordinates are reported in the "AH" column. For example, at chr13:92798976-94371840, Altai and Chagyrskaya have a 3D divergent window identified at chr13:92798976-93847552, while Vindija has a slightly longer window at chr13:92798976-94371840. Table S5 reports the 3D-modifying variants identified in each window.

See supplementary excel file for large tables.

**Table S4: AH-MH 3D divergent windows with less strict thresholds.** In addition to the AH-MH divergent windows characterized in the maintext (Figs. 3A–B) and reported in Tables S2,S3, we report a set of AH-MH windows using less stringent criteria. Instead of requiring all 20 AH-MH comparisons to be more 3D divergent than all MH-MH comparisons, we required the average AH-MH comparison to be more 3D divergent than all MH-MH comparisons. We considered regions in the $75^{\text{th}}$ percentile most diverged using either the mean squared error (MSE) or Spearman-based $(1 - \rho)$ measures. Otherwise, the procedure to identify AH-MH 3D divergent windows is the same as in the Methods. This identifies 252 windows. Although the windows were identified at approximately 1 Mb resolution (Methods), overlapping windows were merged. The "AH" column details for which AH(s) the 3D divergent window was identified (A: Altai, C: Chagyrskaya, D: Denisova, V: Vindija). If a window was identified in two different AHs but they were only partially overlapping, the specific coordinates are reported in the "AH" column. For example, at chr14:69206016-70778880, Altai and Vindija have a 3D divergent window identified at chr14:69206016-70778880, while Chagyrskaya has a slightly shorter window at chr14:69206016-70254592. Table S6 reports the 3D-modifying variants identified in each window.

See supplementary excel file for large tables.

**Table S5: 3D-modifying variants identified inside AH-MH 3D divergent windows.** Each 3D-modifying variant that was identified in an AH-MH 3D divergent window (Fig. 3A–B, Table S3) is reported and described. Columns one through four detail the position (in hg19) and alleles. Column five details for which AH(s) the variant and window was found (A: Altai, C: Chagyrskaya, D: Denisova, V: Vindija). It also provides the 3D divergence score. The format is "AH : *in silico* mutagenesis 3D divergence score : AH-MH 3D-divergent window". For example, chr1:74305804 is a 3D-modifying variant identified in the 1 Mb window chr1:73924608-74973184 in Chagyrksyaka, Altai, and Vindija with a 3D divergence of 0.0279 in *in silico* mutagenesis (Methods). Many 3D-modifying variants are identified in overlapping windows. For example, chr1:159131001 is a 3D-modifying variant identified in both chr1:158859264-159907840 and chr1:158334976-159383552 with 3D divergences of 0.0048 and 0.0049 from *in silico* mutagenesis, respectively. Column six provides the coordinates of the TAD in which the 3D-modifying variant is located and column seven provides the protein coding genes within that TAD. Column eight provides overlap with GTEx eQTL in the format "gene:$P$-value:tissue". Column nine provides overlap with putatively adaptive high-frequency haplotypes from Chen et al. [96]. Column ten provides 1KGP phase 3 allele frequencies by super-population (note: "AFR" includes admixed individuals from the Caribbean and southwestern USA). If alelle frequencies are not present, this variant was not introgressed. Column ten provides a CTCF motif match score for 3D-modifying variants that overlapped CTCF sites defined by Vierstra et al. [108]. For details about all the resources used for these annotations, see the Methods.

See supplementary excel file for large tables.

**Table S6: 3D-modifying variants identified in AH-MH divergent windows with less strict thresholds.** Each 3D-modifying variant that was identified in an AH-MH 3D divergent windows with less strict thresholds is reported. See Table S4 for a list of these windows and the criteria used to identify them. Columns one through four detail the position (in hg19) and alleles. Column five details for which AH(s) the variant and window was found (A: Altai, C: Chagyrskaya, D: Denisova, V: Vindija). It also provides the 3D divergence score and the measure (Spearman-based [spe] or mean squared error [mse]) used to identify the variant. The format is "AH : *in silico* mutagenesis 3D divergence score based on $1 - \rho$ : *in silico* mutagenesis 3D divergence score based on MSE : AH-MH 3D-divergent window". For example, chr1:74305804 is a 3D-modifying variant identified in the window chr1:73924608-74973184 in Chagyrksyaka, Altai, and Vindija by the Spearman-based and MSE measures (with 3D divergence 0.0279 and 0.0015, respectively). This variant is also identified for the overlapping window chr1:73400320-74448896 in Chagyrksyaka, Altai, and Vindija with the Spearman-based measure (0.0063, but not identified with the MSE).

| | | Sequence variability | | 3D genome variability | |
|---|---|---|---|---|---|
| | | marginal P | conditional P | marginal P | conditional P |
| Browning introgressed haplotypes | introgression SHARED across populations | 1.9E-49 | 1.3E-44 | 5.7E-09 | 0.00057 |
| | introgression UNIQUE to one population | 0.039 | 0.019 | 0.14 | 0.066 |
| Browning introgressed haplotypes with Neanderthal filter | introgression SHARED across populations | 1.1E-28 | 3.3E-25 | 1.2E-07 | 0.00047 |
| | introgression UNIQUE to one population | 0.067 | 0.014 | 0.00054 | 0.00013 |
| Vernot introgressed haplotype | introgression SHARED across populations | 0.015 | 0.054 | 0.0015 | 0.005 |
| | introgression UNIQUE to one population | 0.48 | 0.79 | 0.0094 | 0.012 |

**Table S7: Both 3D genome and sequence variability are more important in predicting introgression shared across super-populations than introgression unique to a single super-population.** When considering the relationships between 3D genome variability, sequence variability, and amount of introgression (Supplemental Text, Figs. 5, S13), we consider introgression that was shared across 1KGP super-populations (EAS, EUR, SAS) (white rows) compared to introgression unique to only one super-population (gray rows). We find that 3D genome variability (last two columns) is more strongly predictive of introgression shared among all three super-populations. The analysis was replicated on three sets of introgressed haplotypes. Browning introgressed haplotypes are Sprime segments Browning haplotypes with Neanderthal-matching filter are a subset of the Browning et al. [94] introgressed segments that have 30 putatively introgressed variants that could be compared to the Altai Neanderthal genome and had a match rate of at least 30% to the Altai Neanderthal allele. Vernot haplotypes are S* segments from Vernot et al. [15].

| | | Sequence variability | | 3D genome variability | |
|---|---|---|---|---|---|
| | | marginal P | conditional P | marginal P | conditional P |
| Browning introgressed haplotypes | ALL windows (N = 4749) | 1.90E-49 | 1.30E-44 | 5.70E-09 | 0.00057 |
| | ONLY windows with any evidence of introgression (N = 1950) | 0.0004 | 0.0072 | 1.90E-06 | 3.00E-05 |
| Browning introgressed haplotypes with Neanderthal filter | ALL windows (N = 4749) | 1.10E-28 | 3.30E-25 | 1.20E-07 | 0.00047 |
| | ONLY windows with any evidence of introgression (N = 1604) | 0.042 | 0.19 | 0.0001 | 0.00038 |
| Vernot introgressed haplotype | ALL windows (N = 4749) | 1.50E-02 | 5.40E-02 | 0.0015 | 0.005 |
| | ONLY windows with any evidence of introgression (N = 2657) | 3.40E-05 | 8.40E-07 | 0.00068 | 1.60E-05 |

**Table S8: Compared to sequence variability, 3D variability is a relatively more informative predictor of amount of introgression when considering windows of the genome with any introgression.** When considering the relationships between 3D genome variability, sequence variability, and amount of introgression (Supplemental Text, Figs. 5, S13), we consider a subset of windows with any evidence of introgression (gray rows) compared to all windows (white rows). 3D variability is relatively more informative about the amount of introgression when only considering windows of the genome with any introgressed sequence present (last column). The analysis was replicated on three sets of introgressed haplotypes. Browning introgressed haplotypes are Sprime segments Browning haplotypes with Neanderthal-matching filter are a subset of the Browning et al. [94] introgressed segments that have 30 putatively introgressed variants that could be compared to the Altai Neanderthal genome and had a match rate of at least 30% to the Altai Neanderthal allele. Vernot haplotypes are S* segments from Vernot et al. [15].
